# Characterizing and mitigating confounding by unreplicated evolutionary events in phylogenetic regression

**DOI:** 10.64898/2025.12.04.692436

**Authors:** Qifan Wang, Michael D. Edge, Joshua G. Schraiber, Matt Pennell

## Abstract

When researchers want to test for a direct effect of one attribute on another using interspecific data, they need to account for the confounding influence of other attributes that have phylogenetic structure. Accordingly, a suite of approaches have been developed for this purpose, mostly notably the use of phylogenetic regression. However, it has been shown that such methods can be misled by evolutionary events with large effects. We suggest that one widely-applicable solution to this problem can be found by borrowing both from the statistical genetics literature and by digging into the past of phylogenetic comparative methods. In conventional phylogenetic regression, the phylogenetic variance-covariance matrix is included as a random-effect term; here, we argue that it can be advantageous to also include the leading eigenvectors of this matrix as fixed effects. Building on the recent results of Schraiber et al. (2024), we use mathematical analysis and simulations to show under which scenarios this hybrid strategy is effective. We develop a novel approach for visualizing the contributions of different branches of the phylogeny to the eigenvectors of the variance-covariance matrix. We also illustrate how quantile-quantile plots can be used to assess whether the phylogenetic structure has been effectively controlled. We then apply the mixed approach to investigate the co-evolution of gene expression levels of different genes across the phylogeny of cichlid fishes; we show that including the leading eigenvectors likely reduces the false positive rate. To facilitate the use of our visualization approach and of including eigenvectors in phylogenetic regression models, we present a new R package called EIGER. We argue that the approaches we explore here can both help address the statistical problems that arise from large, historical events and more generally, provide richer insights into the nature of phylogenetic confounding.

## Introduction

It is widely appreciated that correcting for phylogenetic structure is a critical requirement in many, and perhaps most, hypothesis tests involving interspecific data. Indeed, it is so routine in comparative studies that the debates about the validity and purpose of this approach that occurred decades ago (see, for instance, discussions in Harvey et al., 1991) may now seem obscure. The standard conceptual model is that unmeasured traits affect both the response and explanatory variable (i.e., these traits are confounders) and that the unmeasured traits are distributed in a way that reflects phylogeny (Rohlf, 2001, 2006; Uyeda et al., 2018; Westoby et al., 2023). Phylogenetically indepedent contrasts (Felsenstein, 1985), phylogenetic generalized least squares (Grafen, 1989; Martins and Hansen, 1997) or phylogenetic linear mixed models (Lynch, 1991; Housworth et al., 2004) are the workhorse statistical approaches to model this scenario (for simplicity, we will hereafter use the shorthand ‘PGLS’ to refer to these approaches collectively; as we discuss later, the differences are not pertinent to the present study).

However, as pointed out by Uyeda et al. (2018), PGLS, like many other phylogenetic comparative methods (Read and Nee, 1995; Maddison and FitzJohn, 2015), have unacceptably high false-positive rates when the evolution of traits is characterized by large, rare events that are inconsistent with the continuous processes that are assumed by nearly all models. This is unfortunate for two reasons. First, it is correcting for these large events—the type of evolutionary event that define what taxonomists consider a clade—that is often the stated rationale for the use of phylogenetic comparative methods in the first place (Uyeda et al., 2018). Second, large bursts of trait change have been found to be a characteristic dynamic of many phenotypes of interest (Uyeda et al., 2011; Pennell et al., 2014; Landis and Schraiber, 2017).

There have been several suggestions as to what to do about the pernicious influence of unreplicated events in phylogenetic comparative studies. Uyeda et al. (2018) pointed out a conceptually simple approach: if, for example, there had been only one large shift in the mean phenotype over evolutionary time and it was known where on the phylogeny it occurred, one could simply include an indicator variable in the regression model specifying on which side of the shift event a species falls on the phylogeny. In practice, however, this requires a separate method for identifying the shift point (we return to this problem below). Further, it is unclear how best to construct the model if there are multiple mean shifts along the phylogenetic tree. Hidden Markov models have become widely used in phylogenetic comparative studies to mitigate the influence of unmodeled background variation in evolutionary rates in discrete characters (Beaulieu et al., 2013; Caetano et al., 2018); it is not apparent how to translate this idea into the formulation of a generalized regression model. Last, some researchers have explored the use of robust regression techniques (Slater and Pennell, 2014; Adams et al., 2024, 2025); robust regression methods effectively down-weight high-leverage observations, reducing the false positives that result from large-effect, unreplicated events. Although applying robust regression may be effective in some contexts, these approaches cannot guarantee consistent estimation of variance components and lack some of the flexibility of generalized linear models. Moreover, it is unknown how to map the statistical model used in robust regression to any particular evolutionary model, and thus it is hard to gain intuition as to how and why it may work in any given scenario.

Schraiber et al. (2024) showed that both the PGLS model and the linear mixed model (LMM) used in genome-wide association studies (GWAS) can be derived as a special cases of the same general model. This result suggests that tools developed in one field can often be readily applied in the other. This is valuable because GWAS studies are susceptible to the same type of large-effect events as phylogenetic models. In the context of GWAS, researchers are trying to control the confounding influence of population structure (Freedman et al., 2004; Marchini et al., 2004; Helgason et al., 2005; Campbell et al., 2005); as in phylogenetic comparative methods, the LMM used by statistical geneticists includes random effects that are assumed to be distributed in a way that mirrors the relatedness among individuals in the sample (Kang et al., 2008, Kang et al., 2010; Astle and Balding, 2009; Zhou and Stephens, 2012). A fairly common practice in statistical genetics is to use models that include both random effects parameterized by a relatedness matrix and fixed effects for individuals’ coordinates on the eigenvectors of the same relatedness matrix. Although the apparent redundancy of this “belt and suspenders” approach has been recognized in the GWAS literature, it is commonly used and helpful when large environmental effects covary with population structure (Schraiber et al., 2024).

This state of affairs led us to reason that the solution devised by statistical geneticists could be applied to the case of phylogenetic regression: in addition to including the phylogenetic structure as a random effect in the regression, also include several eigenvectors of the phylogenetic variance co-variance matrix (defined below) as fixed-effect terms. In the 1990s, the idea of using phylogenetic eigenvectors was presented as an alternative to the use of mixed models (Diniz-Filho et al., 1998, referred to as Phylogenetic Eigenvector Regression, or PVR); this approach was argued to have inferior statistical properties to PGLS (Rohlf, 2001; Freckleton et al., 2011). We will discuss the reasons for this assessment below. The insight of statistical geneticists is that one could profitably combine these two strategies (i.e., use both LMMs *and* eigenvectors). This mixed strategy was suggested by Schraiber et al. (2024) and used by David et al. (2025) in a recent study to uncover the genetic bases of convergent metabolic innovation across yeasts. To our knowledge, this idea has not been used elsewhere in the context of comparative biology.

We refer readers to our previous paper (Schraiber et al., 2024) for a more complete theoretical analyses of the use of eigenvectors in the context of a PGLS model, but we repeat the key elements of the approach here. In a sample of size *n* tips, we have a predictor, *x* = (*x*_1_, *x*_2_, …, *x*_*n*_)^*T*^ and an outcome variable, *y* = (*y*_1_, *y*_2_, …, *y*_*n*_)^*T*^, both of which are either organismal traits or proxies thereof. To analyze the effect of *x* on *y*, we can use the following regression model:

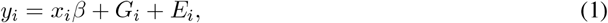

where *β* is the effect of *x* on *y* and *G*_*i*_ and *E*_*i*_ are genetic and environmental components of variance in *y*, respectively. By genetic component, we are referring to features encoded by the genome of a species aside from *x* that might influence *y*. The environmental component refers to non-genetic contributions to the phenotype. In this formulation, the model ignores *G*× *E* interactions, following typical practice. Testing for an effect of *x* on *y* requires accounting for covariance in the genetic component *G*, or else hypothesis tests will not have the desired type-I error rate (Rohlf, 2001, 2006). The *G*_*i*_ values of different species covary owing to shared ancestry, and we can express the expected pattern of covariation with the “phylogenetic variance-covariance matrix”, which will hereafter denote as Σ. Making standard distributional assumptions, the PGLS model can be written as

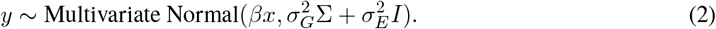

We can modify this model by including eigenvectors of Σ as fixed effects alongside *β*. To do so, we perform the eigenvector decomposition Σ = *V* Λ*V*^*T*^, where *V* = [*v*_1_ *v*_2_ · · · *v*_*n*_] is a matrix containing the eigenvectors of Σ as its columns, and Λ = diag(*λ*_1_, *λ*_2_, …, *λ*_*n*_) is a diagonal matrix with the eigenvalues of Σ on the diagonal. As a variance-covariance matrix, Σ is positive semidefinite, and its eigenvectors can be used to form an orthonormal basis of ℝ^*n*^.

Schraiber et al. (2024) provided a mathematical argument that using the eigenvectors of Σ as fixed effects in a generalized least squares framework could provide the benefits of both approaches: if an eigen-vector of Σ contains some large confounding that is out of proportion with its associated eigenvalue, that confounding can be removed by including the eigenvector in the design matrix. For eigenvectors that are not included in the design matrix, we can still control for spurious association between *x* and *y* caused by the phylogenetic structure by utilizing PGLS. Although Schraiber et al. (2024) showed why this approach could work, it was a small part of an argument targeted elsewhere. Furthermore, they focused primarily on the theoretical issues around how eigenvectors capture different components of the variance from PGLS and not on the performance and use of the approach. Here, we perform a much more extensive set of simulations that evaluate the performance of the method in a wider variety of evolutionary scenarios; develop new mathematical results, and an accompanying visualization technique, that illustrate how the eigenvectors capture phylogenetic structure in specific cases; and illustrate the use of quantile-quantile plots for assessing how effectively researchers have removed the influence of phylogenetic confounding. We also discuss pragmatic issues in the application of this technique, address previously raised critiques of phylogenetic eigenvector regression, and provide a new R package EIGER, that implements the approaches described here.

## Methods

### Visualization of contributions of branches to eigenvectors

Before describing the method of visualizing contributions of branches to eigenvectors, we make one note of clarification: our approach is distinct from the computation of “phylogenetic principal components analysis (pPCA)” (Revell, 2009; Polly et al., 2013). In that case, one is computing the eigenvectors of a matrix of traits *X* and using the Σ matrix to remove the phylogenetic signal. This is mathematically and conceptually distinct from what we are doing here and is used to solve a different problem (see Uyeda et al., 2015; Bookstein, 2019).

To visualize how eigenvectors capture the phylogenetic structure, we developed a tool to compute the contributions of specific branches of the species tree to the eigenvectors of the phylogenetic variance-covariance matrix (Figure 1A). Population and statistical geneticists have developed the idea of the expected genetic relatedness matrix given a gene-genealogical tree relating haplotypes and assumptions about the mutation process (McVean, 2009; Fan et al., 2022; Zhang et al., 2023; Lehmann et al., 2025), variously called an expected genetic-relatedness-matrix (eGRM), an ancestral-recombination-graph GRM, or a branch GRM; we use the term eGRM to refer to all of these approaches. These eGRMs can be expressed as a weighted sum of eGRMs corresponding to each branch of the tree (Fan et al., 2022). Although the eGRM approach was designed in the context of genealogical trees, the principle applies to phylogenetic trees just as well. Here we extend this idea to our Σ matrix, assuming that traits evolve following a Brownian motion (BM) model across the phylogenetic tree. We denote this particular version of Σ as *C*. Given the phylogenetic tree T and the BM diffusion rate of *σ*^2^, and the contribution of a branch *e* to *C* is

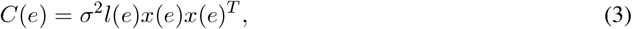

where *l*(*e*) is the branch length, and *x*(*e*) is a binary vector for the descendancy relationship, with *x*(*e*)[*i*] being 1 if the *i*-the tip is a descendant of branch *e* and 0 otherwise. Then, *x*(*e*)*x*(*e*)^*T*^ is a matrix where the *i, j* entry is 1 if both species *i* and *j* descend from *e*. We can compute *C* for the entire tree T by summing the contributions from all branches:

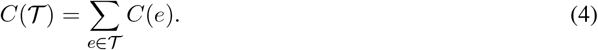

**Fig 1:**
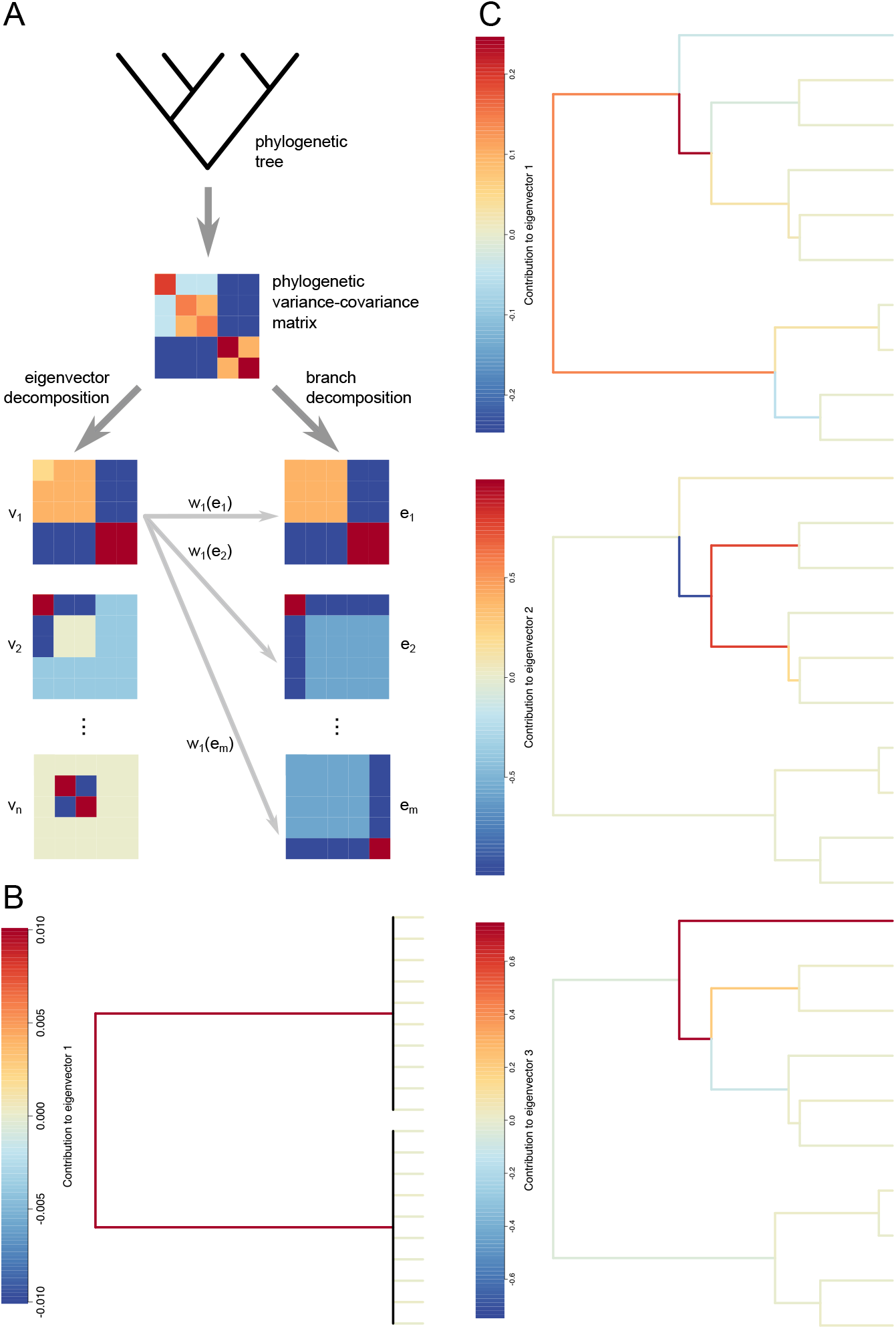
Contributions of branches to the eigenvectors of a phylogenetic variance-covariance matrix Σ. (A) Flowchart showing the method to calculate the contributions of branches to the eigenvectors of Σ. (B) Contributions of branches to *v*_1_ of the double-centered phylogenetic variance-covariance matrix 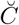 of a phylogenetic tree depicting Felsenstein’s Worst Case (FWC). Each clade contains 10 species. The vertical branches are black because the tip branches slightly differ in the their contributions to *v*_1_. (C) Contributions of branches to the first three eigenvectors of 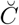 of a Yule tree containing 10 species.

Note that this equation will result in exactly the variance-covariance matrix in the standard Brownian motion model: the *i, j* entry will be the sum of branch length share by species *i* and *j* scaled by the rate of Brownian motion, *σ*^2^.

The next step would be to perform an eigenvector decomposition. Although we can decompose *C* itself, in principle, it would be best practice to first perform a double-centering transformation. This transformation can be interpreted as unrooting the tree so that the contributions coming from the two branches stemming from the root would merge into one. This is helpful for visualization, since the partition of the species into descendants and non-descendants of the two branches would be the same: if a species is a descendant of one branch, than it is a non-descendant for the other. This double-centering transformation also helps build the connections between our methods and many previous methods using PCs for controlling confounding in both statistical genetics and phylogenetics contexts (Diniz-Filho et al., 1998; McVean, 2009; de Vienne et al., 2011). We can show that the eigenvectors of the double-centered version of the matrix, which we denote as 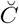, are exactly the same as those of the double-centered distance matrix 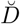 used in Diniz-Filho et al. (1998), with eigenvalues differing by a constant (see Section 3 of the supplementary material). Following de Vienne et al. (2011), we can also show that the eigenvectors of 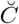 correspond to the projections of the species onto the primary axes of a PCA, if we consider the *n* tips of the phylogenetic tree as *n* points in R^*n*^ space (see Section 4 of the supplementary material). Due to the convenient properties of double-centeredness, the usage of a double-centered matrix is even more prevalent in statistical genetics, where the data are essentially always mean-centered column-wise before being used to construct a GRM (see Yang et al., 2010) or eGRM (see Fan et al., 2022), which are statistical genetics equivalents of *C*. However, we’d also like to note that even though it is theoretically preferable to use the double-centered version 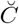 to get eigenvectors, in practice using a non-centered *C* would not change the results by much when doing phylogenetic regressions (see Section 5 of the supplementary material for theoretical analyses and Fig. S4, S5, and S14 for simulation results). If we perform a double-centering transformation on *C*, then we obtain:

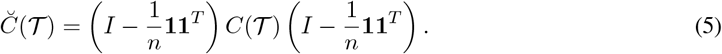

The eigenvector decomposition of 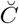 is:

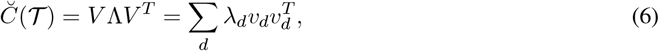

where the summand 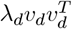 in the Eq. 6 refers to the contribution from one of the orthogonal dimensions to 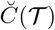. Finally, we can obtain the contributions of branches to the eigenvectors by running a linear regression:

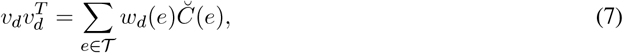

where *w*_*d*_(*e*) is the contribution of branch *e* to the *d*-th eigenvector of the phylogenetic tree. Note that *w*_*d*_(*e*) does not necessarily have to be positive, and we only care about the relative weights across the branches. Thus *λ*_*d*_ does not affect the relative weights of Σ(*e*) here: including *λ*_*d*_ into the equation would multiply the weights *w*(*e*) by a constant factor of *λ*_*d*_. Once the contributions of branches to each of the eigenvectors are obtained, one can also compute the overall contributions of branches, which can then be used as a metric for the importance of the branches (see Section 6 of the supplementary material).

### Software implementation

We have developed a new open-source R package `EIGER` that includes two key capabilities. First, using the functions `compute_branch_contributions()` and `plot_branch_contributions()`, researchers can evaluate the contributions of different branches on a phylogenetic tree to its eigenvectors. Second, we have built a wrapper for the core function of phylolm (Ho et al., 2016) so that researchers can include eigenvectors as fixed effects in the general PGLS model. This package is available on GitHub https://github.com/applied-phylo-lab/eiger.

### Simulation analysis

Schraiber et al. (2024) investigated the performance of using eigenvectors in PGLS in simple scenarios. In particular, they studied Felsenstein’s Worst Case (hereafter FWC), a scenario developed by Felsenstein (1985) and examined in more detail in Uyeda et al. (2018), in which the only phylogenetic structure in a dataset is between two clades, and there is a shift in the mean trait values along a branch leading to one of the clades; additionally they considered the case of BM evolved on a Yule tree. Here, we are exploring a much larger tree space by considering different tree topologies of varying sizes. To test the effect of tree size, we simulated Yule trees, caterpillar trees, fully balanced trees, and coalescent trees of 8, 16, 32, 64, 128, 256, 512, and 1024 species (Fig. 2). For each combination of tree type and tree size, we generated 500 different ultrametric trees with varying branch lengths. Then we simulated *x* and *y* as arising independently from a Brownian motion along each of the trees. For simulations, we used 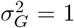 and 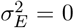. Although real datasets feature non-trivial contributions from environmental noise (for further details, see Hansen and Bartoszek, 2012; Silvestro et al., 2015; Pennell et al., 2015), the results would be qualitatively similar (Schraiber et al., 2024). After *x* and *y* were generated, we compared the performance of using different numbers of eigenvectors in OLS and PGLS for phylogenetic regression using the functions `lm()` and `phylolm()`in the `phylolm R` package (Ho and Ané, 2014; Ho et al., 2016). In Section 7 of the supplementary material, we show that our results apply equally to PGLS and PGLMM formulations of phylogenetic regression.

**Fig 2:**
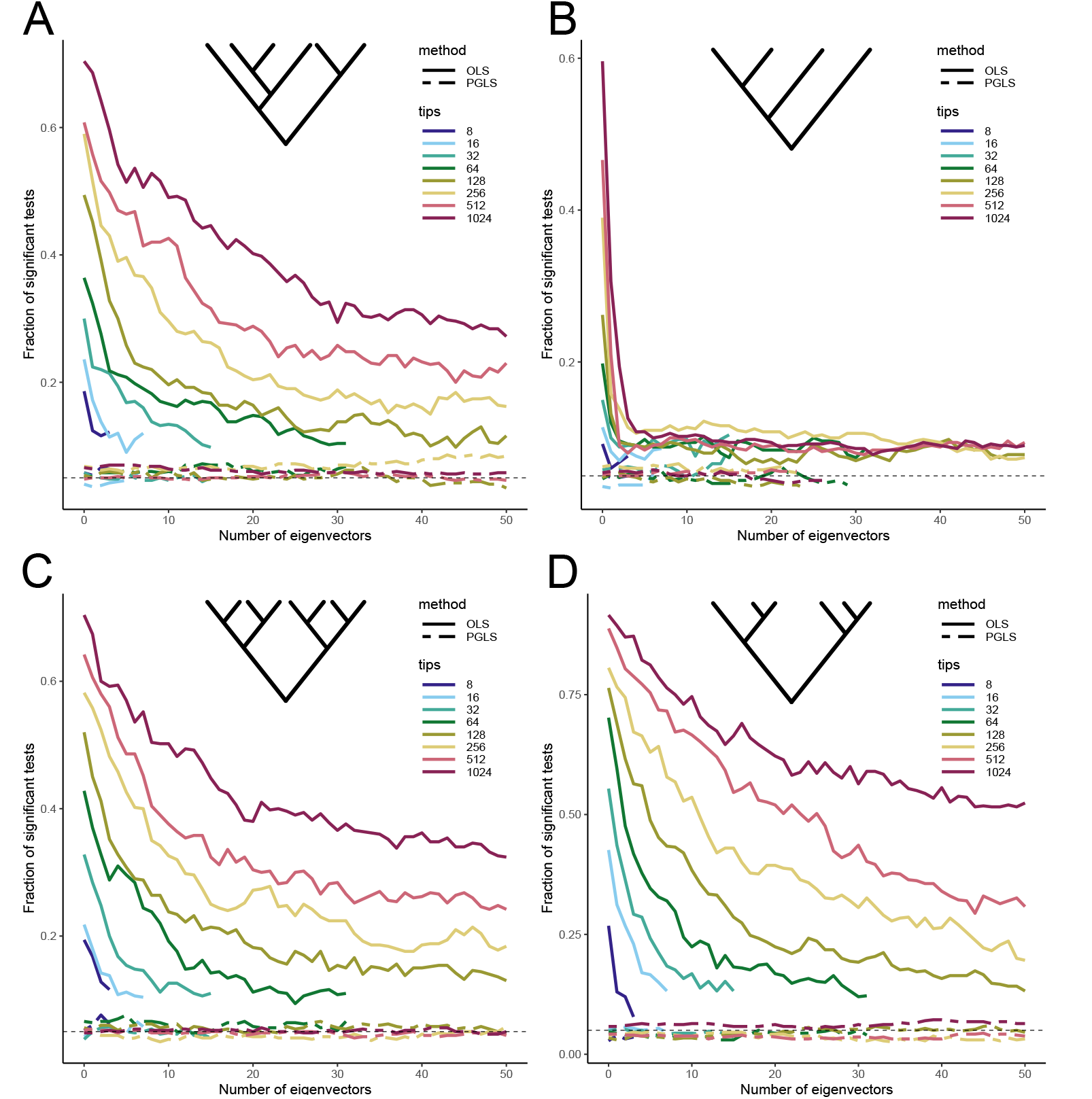
The false positive rate after adding eigenvectors in OLS and PGLS for different types of trees when the data is generated by a pure Brownian motion with no additional shifts. (A) Performance of OLS and PGLS in models related by a Yule tree of varying numbers of species. For simplicity, a Yule tree with only 6 tips is shown. The horizontal axis shows the number of eigenvectors included as covariates, and the vertical axis shows the fraction of tests that would be significant at the 0.05 level. (B) Performance of OLS and PGLS in models related by a caterpillar tree of varying numbers of species. For simplicity, a caterpillar tree with only 4 tips is shown. The horizontal axis shows the number of eigenvectors included as covariates, and the vertical axis shows the fraction of tests that would be significant at the 0.05 level. (Note that when a very large number of eigenvectors were included, the PGLS model could not be estimated.) (C) Performance of OLS and PGLS in models related by a fully balanced tree of varying numbers of species. For simplicity, a fully balanced tree with only 8 tips is shown. The horizontal axis shows the number of eigenvectors included as covariates, and the vertical axis shows the fraction of tests that would be significant at the 0.05 level. (D) Performance of OLS and PGLS in models related by a coalescent tree of varying numbers of species. For simplicity, a coalescent tree with only 6 tips is shown. The horizontal axis shows the number of eigenvectors included as covariates, and the vertical axis shows the fraction of tests that would be significant at the 0.05 level.

We then explored the performance of the methods when there are non-Brownian shifts in trait value on the trees, generalizing the work of Schraiber et al. (2024), who examined FWC with one shift on one of the clades. Here, we revisited the same scenario with a quantile-quantile plot (QQ plot) of *p*-values. If the null hypothesis is true (i.e., there is no association between traits), the *p*-values will follow the expected distribution and they will fall on the 1:1 line of the QQ plot; by convention, we use −log_10_(*p*), which will be uniformly distributed ∈ [0, 1]. If there are some −log_10_(*p*) that are bigger than expected, this may indicate a true signal of an association; if the entire distribution of −log_10_(*p*) is inflated, this likely means that the phylogenetic structure is confounding the tests. Although this is a routine procedure in statistical genetics (in a GWAS, there will always be many tests performed), we are only aware of one application of this idea to phylogenetically structured data (Hale et al., 2025).

For FWC with one shift on one of the clades, we ran a total of 100 replicates and created QQ plots for four methods: OLS, OLS with eigenvector *v*_1_, PGLS, and PGLS with *v*_1_. (Fig. 3A) We applied the same tool to a slightly more complicated scenario, where there is a non-Brownian shift on a branch of a Yule tree (Fig. 3B). We generated a Yule tree of 100 species and introduced a shift on one of the branches. We simulated *x* and *y* as arising from independent Brownian motions along the tree, and we generated the shift by adding a normal random variable with a large standard deviation (*s* = 100) to the species that are descendants of the branch where the shift occurs. We ran a total of 100 replicates and we created QQ plots for OLS and PGLS with different numbers of eigenvectors (from 0 up to 30).

**Fig 3:**
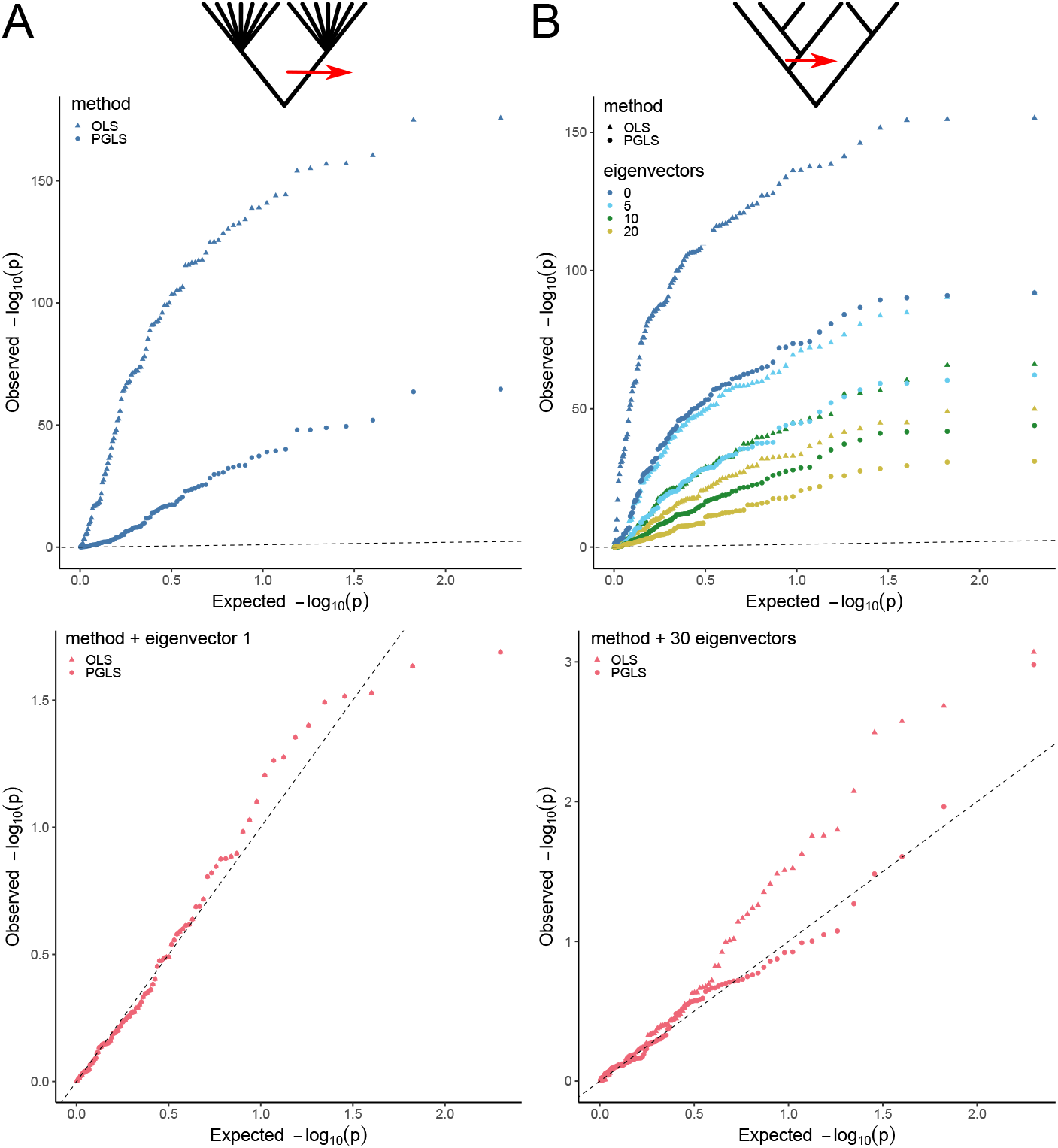
Quantile-quantile plots (QQ plots) showing the performance of different methods in controlling phylogenetic confounding. (A) QQ plots showing the performance of different methods in controlling confounding in Felsenstein’s worse case (FWC) with a non-Brownian shift in both predictor and outcome variables. For simplicity, only 7 tips are shown for each clade; in simulations, each clade contains 100 tips. The red arrow represents the non-Brownian shift on the branch giving rise to one of the two clades. The dotted line shows the case when the observed *p*-values exactly match the expected null *p*-values. The top panel shows the case when the methods fail to control the confounding, and the bottom panel shows the case when the methods manage to control the confounding. The dots for OLS + eigenvector *v*_1_ and PGLS + *v*_1_ are completely over-lapping. (B) QQ plots showing the performance of different methods in controlling confounding in model related by a Yule tree with a non-Brownian shift in both predictor and outcome variables. For simplicity, only 6 tips are shown; in simulations, the Yule tree contains 100 tips. The red arrow represents the non-Brownian shift on one of the branches. The dotted line shows the case when the observed *p*-values exactly match the expected null *p*-values. The top panel shows the case when the methods fail to control the confounding, and the bottom panel shows the case when the methods manage to control the confounding.

Building on the last scenario, we are interested in how many eigenvectors we need to include as covariates in the model to mitigate the confounding when there is a non-Brownian shift of different sizes on different branches of the tree. We generated a Yule tree with 16 tips, and we simulated *x* and *y* arising independently from a Brownian motion along each of the trees. Then we selected a branch where the shift occurs, and we considered two different types of shifts. The first type is the same as in the previous simulations, where the shift is generated by adding a normal random variable with a varying standard deviation to *x* and *y* values of the species that are descendants of the branch where the shift occurs. Biologically, this can be interpreted as an acceleration of the evolutionary rate on that particular branch where the shift occurs (*sensu* Landis and Schraiber, 2017). The second type of shift is generated by adding a constant of a varying size to *x* and *y* values of the species that are descendants of the branch where the shift occurs. A shift of this type can be interpreted as a shift in the optimal trait value within the corresponding clade. As in the previous simulations with no shifts, we compared the performance of using different numbers of eigenvectors in OLS and PGLS for phylogenetic regression. For each combination of shift position, shift type, and shift size, we ran a total of 100 replicates (Fig. 4).

**Fig 4:**
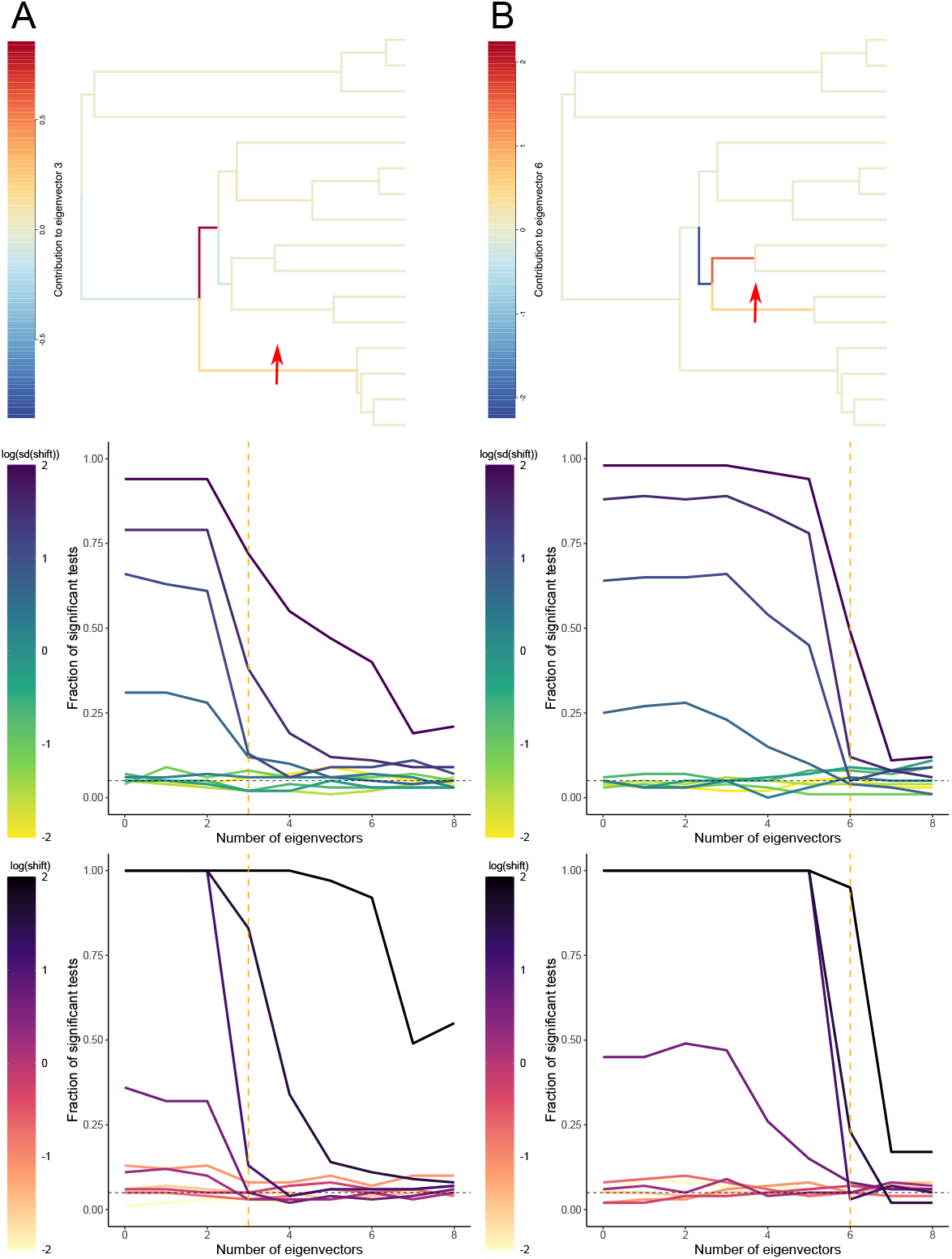
The performance of using eigenvectors in phylogenetic generalized least squares (PGLS) to remove phylogenetic confounding in a model with 16 tips related by a Yule tree with non-Brownian shifts in both predictor and outcome variables. The top panels show the contribution of branches to one eigenvector of the variance-covariance matrix *C*. The red arrows indicate the branches where the shifts occur. The middle panels show the performance of using eigenvectors in PGLS to remove phylogenetic confounding in the model, where the shift is simulated by adding a normal random variable with varying standard deviation to the species that are descendants of the branch. The horizontal axis shows the number of eigenvectors included as covariates, and the vertical axis shows the fraction of tests that would be significant at the 0.05 level. The vertical orange dashed lines show the number of eigenvectors corresponding to the dimension in the top panels. The bottom panels show the performance of using eigenvectors in PGLS to remove phylogenetic confounding in the model, where the shift is simulated by adding a constant of varying size to the species that are descendants of the branch. The horizontal axis shows the number of eigenvectors included as covariates, and the vertical axis shows the fraction of tests that would be significant at the 0.05 level. The vertical orange dashed lines show the number of eigenvectors corresponding to the dimension in the top and middle panels. (A) The shift falls on a branch that contributes the most to *v*_3_ of *C* among all the first 8 eigenvectors. (B) The shift falls on a branch that contributes the most to *v*_6_ of *C* among all the first 8 eigenvectors.

### Application to gene expression data from cichlids

As an illustration, we used data on gene expression from across a radiation of African rift lake cichlids from El Taher et al. (2021). Though not the original purpose of the study, we followed the study design of Cope et al. (2020) in testing whether pairs of genes showed evidence of co-evolution in their mRNA counts within the same tissue. Using the approach described above, we visualized how the first three eigenvectors capture the phylogenetic structure of the 73 species involved (Fig. 5A). We then selected 1000 random pairs of protein-coding genes that have non-zero gene expressions in lower pharyngeal jaw bones across all the species, and we performed PGLS with eigenvectors to see how some correlations can be spurious due to uncorrected phylogenetic confounding (Fig. 5B). Finally, we created QQ plots for the tests on these 1000 pairs with varying numbers of eigenvectors included as fixed effects (Fig. 5C).

**Fig 5:**
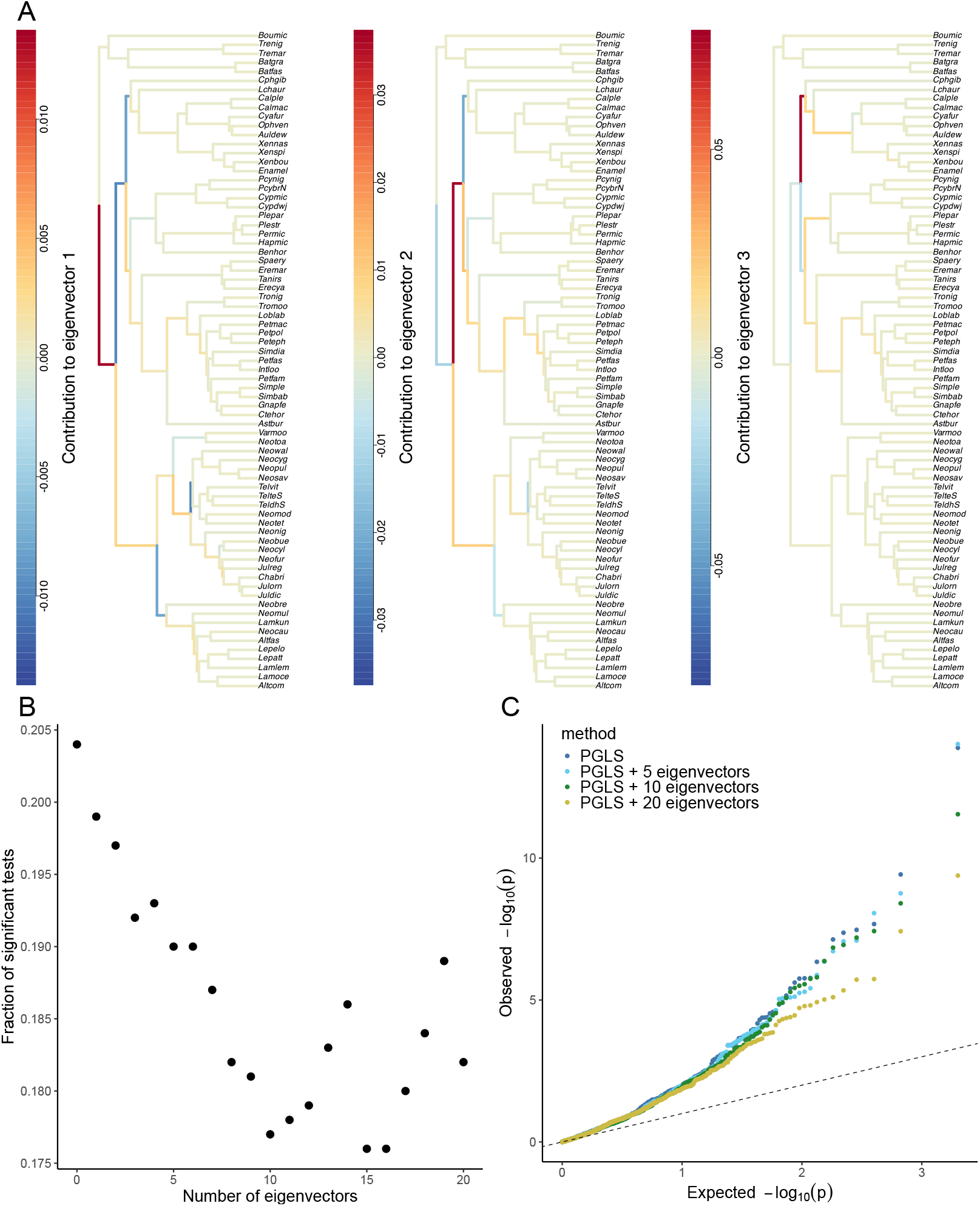
Effect of including phylogenetic eigenvectors on detection of coevolution of gene-expression levels in African cichlids. (A) Contributions of the branches to the first 3 eigenvectors of the phylogenetic variance-covariance matrix *C* of the phylogenetic tree for the 73 cichlid species. (B) The overall number of significant pairs first decreases and then fluctuates as more eigenvectors are included in PGLS. The horizontal axis indicates the number of eigenvectors included as fixed effects, and the vertical axis shows the proportion of significant pairs out of 1000 randomly selected pairs. (C) QQ plot showing the performance of using different numbers of eigenvectors included as fixed effects in PGLS. The dotted line shows the case when the observed *p*-values exactly match the expected null *p*-values.

## Results

### Description of simulation results

To examine the visualization tool we developed, we first checked how the branches contribute to the first eigenvector *v*_1_ of 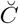 in FWC. Since the only phylogenetic structure in this scenario is the split between the two clades, we expect *v*_1_ to capture this split. This is exactly what we observe in Fig. 1B, as the two branches from the root splitting the two clades contributes the most to *v*_1_, while all the other branches have nearly no contributions.

We also looked at the contributions of branches to the first three eigenvectors of 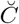 on a Yule tree of 10 species. As shown in Fig. 1C, *v*_1_ captures the largest split between the two largest clades as well as some splits within these clades. *v*_2_ captures the split between the two-species clade and three-species clade within the large six-species clade, finer and more recent phylogenetic structure than in *v*_1_. *v*_3_ explains the split of the outgroup from the rest of the five species in the large 6-species clade. More generally, the dimension of the largest variance among all the species would generally correspond to the splits or branches that give rise to the most even partition of the species. The visualization method can also be easily extended to realizations of Σ matrix other than 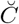 (see Figs. S1, S2, and S3). In Section 5 of the Supplementary Material, we explore the impact of centering on our visualization of the eigenvectors.

Fig. 2 shows the false positive rate after adding eigenvectors in OLS and PGLS for different types of trees when the data is generated by a pure Brownian motion with no additional shifts. In general, we see that using eigenvectors in conjunction with a random effects term capturing the phylogenetic structure (i.e., with PGLS) performs much better than using eigenvectors without the random effects term (i.e., with OLS); recall that using eigenvectors with OLS is essentially identical to the Phylogenetic Eigenvector Regression approach developed by Diniz-Filho et al. (1998), which is known to underperform PGLS under a purely Brownian model (Rohlf, 2001). As we have discussed, including eigenvectors of *C* as fixed effects can correct for punctate effects that are specific to a branch or region of the tree, whereas including *C* as the covariance structure in PGLS does a better job in correcting for phylogenetic confounding that is diffuse across the tree. In our simulations, the trees generally have complex structure, and thus including the eigenvectors in OLS ignores some of the later dimensions and fails to correct for diffuse confounding that occurs due to recent splits and other more subtle features of the phylogeny. We also generally observe that for OLS, adding more eigenvectors decreases the false positive rates, while adding more eigenvectors has little effect on the false positive rates when using PGLS. As later eigenvectors capture finer-scale structure of the phylogeny, including more eigenvectors in OLS can indeed correct for more of the smaller phylogenetic confounding. In contrast, in PGLS, the diffuse structure of the phylogeny is already taken into account by including *C* in the random effects term, and thus additional eigenvectors are redundant. Another clear pattern is that for larger trees (with more species), the false positive rate of using the same number of eigenvectors in OLS also tends to be larger, meaning that one would need to include more eigenvectors in OLS to correct for phylogenetic confounding. Intuitively, this is because the same number of eigenvectors capture a smaller proportion of the tree in a larger tree compared to a smaller tree. Analyses on non-ultrametric trees also generated similar results (Fig. S6).

Among the four types of trees we simulated, caterpillar trees stand out as the false positive rates of using eigenvectors in OLS drop much more quickly compared to other types of trees (Fig. 2B). This is because caterpillar trees have relatively simple structures, and when the internal branches are short, part of the caterpillar trees can be approximated as a star tree, thus collapsing into something similar to what we observe in FWC: even though there are multiple species, there is very little structure and thus can be captured by the first few eigenvectors. Interestingly, for OLS, after having 5 to 6 eigenvectors, adding more eigenvectors does not further decrease the false positive rate by much, and it remains an open question how the false positive rate curve would behave as the number of eigenvectors included as fixed effects approaches the number of species in the tree — when too many eigenvectors are included, the OLS model simply cannot be estimated because there are too many parameters to estimate compared with the number of observations.

Although false positive rates provide us with a summary of the results of multiple replicate tests, we also lose information by collapsing a list of values into one. Here we show how QQ plots of *p*-values can help us judge whether the confounding effect of phylogenetic structure is plausibly controlled. Expanding on the simulations of FWC with one shift in Schraiber et al. (2024), we created QQ plots for 100 replicates on a tree composed of two clades, each containing 100 species. As the first eigenvector *v*_1_ can capture the split of the two clades, we expected both OLS and PGLS with the first eigenvector to be able to remove the phylogenetic confounding. This is exactly what we observe in Fig. 3A, where both methods that include *v*_1_ have the observed *p*-values aligning with the expected *p*-values (bottom panel), whereas both methods that do not include *v*_1_ as fixed effect give rise to a large number of much smaller *p*-values than expected (top panel). It is also noteworthy that when including *v*_1_ as covariate, the results of OLS and PGLS overlap exactly, which agrees with the results from Schraiber et al. (2024). We can mathematically explain this result by applying equations (16) and (18) in Schraiber et al. (2024) to FWC (see Section 8 of the supplementary material).

We also created QQ plots for cases where there is a shift on the 100-tip Yule tree (Fig. 3B). Similar to previous simulations, the group of methods using PGLS in general perform better than their OLS counterparts, and for OLS, as more eigenvectors are included, the observed distribution of *p*-values get closer to that expected under the null hypothesis. However, because there is now a non-Brownian shift in the data, we see that additional eigenvectors helps to mitigate the confounding. As the top panel shows, even 20 eigenvectors fail to fully correct the confounding in the phylogenetic regression; the bottom panel shows that we need to include as many as 30 eigenvectors in combination with PGLS to control the false positive rate. On the other hand, even with 30 eigenvectors, OLS fails to generate uniform *p*-values. This illustrates that including eigenvectors as covariates complements PGLS. We have a Yule tree with lots of diffuse phylogenetic structure, so we need PGLS to correct for the overall confounding. On the other hand, we also have a large shift on one of the branches, which leads to confounding that’s out of scale with the tree and requires eigenvectors as covariates.

However, from the previous case it is not immediately clear why we would need 30 eigenvectors to fully correct for the confounding caused by the shift. Therefore, we turned to a much smaller tree containing just 16 species and investigated how the size of the shift and where it occurs on the tree impact the number of eigenvectors we need to include in phylogenetic regressions to fully mitigate the confounding. We considered two types of shifts: one corresponding to an acceleration of the evolutionary rate on the branch, and the other corresponding to a change in the optimal trait value for the clade. We focus our discussion on two representative scenarios (for more scenarios, see Figs. S7, S8, and S9). In Fig. 4A, the shift occurs on the branch that leads to the four-species clade at the bottom of the tree, whereas in Fig. 4B, the shift occurs on a more recent branch that gives rise to only two species. Across both scenarios and both types of shifts, a common pattern is that the false positive rate increases with the shift size but decreases with the number of eigenvectors included as fixed effects. What is more noteworthy, however, is that the false positive rate drops sharply in most cases once we include *v*_3_ in Fig. 4A and *v*_6_ in Fig. 4B. Using the tool introduced at the beginning to visualize the contributions of branches to the eigenvectors of the variance-covariance matrix, we see that among the first eight eigenvectors, the branch where the shift occurs in Fig. 4A contributes mostly to *v*_3_, and in the case of Fig. 4B, the branch contributes mostly to *v*_6_. This explains why including *v*_3_ and *v*_6_ in the two cases would be especially helpful, since these eigenvectors capture the branches where the shifts occur. Comparing the middle and bottom panels, which correspond to the two types of shifts we simulated, we observe that the results are qualitatively similar for both types of shifts. The combined strategy is slightly more vulnerable to cases when there is a large shift on the optimal trait value, since this type of shift was generated by adding a large constant to the species that are descendants of the branch where the shift occurs. The other type of shift, corresponding to an acceleration of evolutionary rate, was generated by adding a random normal variable with some large standard deviation, so across replicates, the effect of the shift is smaller on average.

In empirical data analyses, it is often unclear whether there is a non-Brownian shift on the phylogenetic tree, and if there is, where the shift occurs. Therefore, we revisited the same case above from an alternative perspective — how the false positive rate can be impacted by where the shift falls on the tree if no eigenvectors are included at all in PGLS (Fig. S10). We utilized the visualization tool for the contributions of branches to the eigenvectors of the variance-covariance matrix and computed the overall contributions of the branches (see Section 6 of the supplementary material), which we used to rank the branches. In general, when the shift falls on a branch that has a higher ranking (i.e., one that makes a large overall contribution), the false positive rate of PGLS alone is higher. This implies that the branches making the most overall contributions are most likely to create large confounding in phylogenetic regressions. However, branches that make less overall contributions can also create large confounding when the shift size is large.

We further looked at cases with multiple shifts on the tree (Figs. S11, S12, and S13). In general, we observe the same pattern as in the case of a single shift on the tree: if we include all the leading eigenvectors as fixed effects, then the minimum number of eigenvectors we need to include is equal to the last dimension to which the branches where the shifts occur make large contributions.

Finally, we investigated cases when *β*, the true effect of *x* on *y* is non-zero (Figs. S15, S16, and S17). In general, our combined method tends to lose power if too many eigenvectors are included as fixed effects. This is especially true when the tree size and the effect size *β* are small (Fig. S15). When the tree size is small, as is in cases of sparse sampling, it is expected that tests will have lower power; when the effect size *β* is small, the value would be close to zero and hard to detect unless more data (number of species) are included. We then looked at cases when there is also a shift on the tree, and our combined method has the highest power when the shift size and *β* are large, corresponding to cases that deviate most from the null scenario (Figs. S16 and S17).

### Empirical example

We first applied our visualization tool to see how the eigenvectors capture the structure of the phylogenetic tree of the 73 species of African rift lake cichlid fishes (Fig. 5A). Similar to our simulations, *v*_1_ and *v*2 capture the splits between large and medium clades, whereas *v*_3_ captures the structure on finer structures and more recent splits.

We then ran PGLS with varying numbers of eigenvectors to test the number of pairs of genes that have significant correlations. As is shown in Fig. 5B, when no eigenvectors are included as fixed effects, around 20% of the 1000 randomly selected pairs show significant correlations. As we include more eigenvectors, this fraction keeps decreasing until around 10 eigenvectors. When even more eigenvectors are included, the fraction of significant tests starts to fluctuate between 17.5% and 19%. The initial drop in the fraction of significant tests suggests that some of the correlations can be spurious if we fail to use the first few eigenvectors to remove the large confounding effect of the tree. The later fluctuation of the fraction suggests that adding more eigenvectors might not be necessarily better; after correcting for the large confounding effects using the first few eigenvectors, adding more eigenvectors might do little to remove additional confounding and will reduce power.

Finally, we created a QQ plot for the tests involving these 1000 pairs of genes (Fig. 5C). A key difference between the QQ plot here and the QQ plots in Fig. 3 is that in Fig. 3, we know that *x* and *y* evolve independently, and thus we know the true distribution of *p*-values, whereas here we do not know in reality how many of the 1000 randomly selected pairs indeed have significant correlations. In a statistical genetics context, it has been argued that if most traits are highly polygenic (or even omnigenic, Boyle et al., 2017), then we expect a general inflation of *p*-values (Yang et al., 2011). Therefore, we can only use the null distribution, a uniform distribution, as expected *p*-values, to test whether there are more significant pairs than the 5% threshold. As Fig. 5C shows, there are indeed quite a lot of pairs that have significant correlations across all the methods, but adding more eigenvectors does help us remove at least part of the phylogenetic confounding and reduce false positive rates.

## Discussion

In this paper, we have advanced the argument that, under some biologically realistic scenarios, a mixed strategy of including eigenvectors of the phylogenetic variance-covariance matrix as fixed effects and using the same matrix to model correlated residuals/random effects performs better than either approach on its own. This may be a counterintuitive proposal. These approaches were previously seen as competing alternatives when they were first developed in phylogenetic comparative methods (Rohlf, 2001; Freckleton et al., 2011). And at first glance, it seems like such a mixed strategy is essentially “double-counting” the phylogeny as it appears twice in the regression equation. But as we argue here and elsewhere (Schraiber et al., 2024), comparative biologists can profit from having seen the evolution of parallel developments in statistical genetics. Statistical geneticists re-discovered the use of principal components to control for ancestry (Zhang et al., 2003; Price et al., 2006), and then subsequently re-discovered the idea of using a LMM for the same purpose (Yu et al., 2006; Kang et al., 2008) — although in the latter case, both comparative biologists and statistical geneticists re-invented an approach developed by quantitative geneticists decades earlier (Henderson, 1984). Statistical geneticists subsequently had the insight that these two could be combined, which to our knowledge had not been considered in comparative biology. The mixed strategy appeared to work so well in that context for much the same reasons we have documented here: disproportionally large effects are not captured well by standard LMM/PGLS approaches and that including the eigenvectors can therefore rescue the statistical model (Price et al., 2006; Tucker et al., 2014) — this was demonstrated mathematically by Schraiber et al. (2024). We emphasize that our results also are in agreement with previous assessments of the use of PVR (Rohlf, 2001; Laurin, 2010; Freckleton et al., 2011; Adams and Church, 2011; Chen and Niu, 2024); except in corner cases such as FWC, *only* including eigenvectors in an OLS regression is a suboptimal strategy. In that case, as our visualization tool shows, the two branches from the root make the largest contribution to the eigenvectors with non-zero eigenvalues and hence also the most overall contributions.

Much of the debate surrounding the PVR approach, both by supporters and critics, was on how many eigenvectors to include in the analysis (Adams and Church, 2011; Freckleton et al., 2011; Diniz-Filho et al., 2012a,Diniz-Filho et al., 2012b). If we do not know where on the tree shifts have occurred, it remains an open problem as to know in advance how many eigenvectors to include; but we emphasize that this is still very much an open problem in statistical genetics as well. Adding eigenvectors may mitigate confounding but will come at a loss of power. Here we point out that if one is doing many phylogenetic tests, for example if one is using comparative data to find associations between traits and genomic features (often referred to as ‘PhyloG2P’ tests Smith et al., 2020), QQ plots are one way to assess whether the confounding has been removed but this is not a sure fire guide as the null distribution of *p*-values will depend on the genetic architecture of the traits (Yang et al., 2011). Nonetheless statistical geneticists have gained rough intuition of what might be an appropriate number in a given scenario and we see no reason why phylogenetic biologists would not do the same. Last, we emphasize that the approaches presented here can be used in comparative studies even if one does not include eigenvectors in the model. For example, one could map the contribution of different branches to the leading eigenvectors and either include that branch as an indicator variable (following the suggestion of Uyeda et al., 2018) or else assess whether one’s analysis is potentially susceptible to being misled by unobserved shifts on these key branches.

Although phylogenetic biologists can certainly learn a lot from statistical geneticists, we end by emphasizing that this is a two-way street. Indeed, as we mentioned above, it is now apparent that there was a missed opportunity for statistical geneticists to take inspiration from PVR and PGLS from phylogenetics rather than re-inventing it themselves years later (for a recent example of a human genetics method that draws inspiration from PGLS, see Patel et al., 2025). The visualization technique we have developed here, which extends theoretical developments in statistical genetics (Fan et al., 2022), could provide new insights into how exactly population structure confounds trait-mapping and related problems and how to mitigate its effects. Looking forward, as tree-thinking becomes increasingly central to the theory and practice of statistical and population genetics (Lewanski et al., 2024; Nielsen et al., 2025), we anticipate that methods developed in phylogenetic comparative biology will become even more pertinent to these fields.

## Data Availability

Code used to generate the results in this paper can be found at https://github.com/applied-phylo-lab/Phylo_PC.

## Acknowledgments

This was supported by startup funds from Cornell University and by NIGMS awards to MP (R35GM151348) and MDG (R35GM137758). We thank Ed Bucker, Jason Mezey, and the Pennell and Buckler labs for critical comments on this work.

## Supplementary material

### 1 Contributions of each branch to the eigenvectors of eGRM

In statistical geneticists, the equivalent of the phylogenetic variance-covariance matrix *C* would be the genetic relatedness matrix [GRM, see 9] or its expected version [sometimes called an eGRM, see 3], both of which are used to capture genetic relatedness between individuals. The data used to generate these two matrices are usually some *n* × *m* genotype matrix *G*, where each of the *n* rows represents an individual, and each of the *m* columns represents a site (or a SNP). The calculation of GRM is directly based on *G*, whereas the calculation of eGRM is usually based on some tree or ARG (ancestral recombination graph) inferred from *G*. Both GRM and eGRM are collapsed over the number of sites and have a dimension of *n* × *n*. Below we briefly show the key steps of calculating an eGRM from a tree 𝒯 that is inferred from *G*.

Let *x*(*e*) be the binary vector for a branch *e* ∈ 𝒯, where *x*(*e*)[*i*] is 1 if the *i*-th tip is a descendant of branch *e* and 0 otherwise. Let *l*(*e*) be the length of branch *e*. Then we can get the eGRM of the branch *e*:

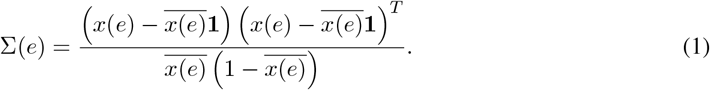

The eGRM of the entire tree would be a weighted sum of branch eGRMs:

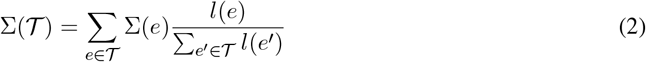

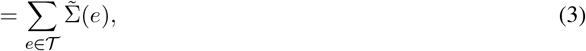

where 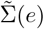 is a weighted eGRM representing the contribution of branch *e* to the tree eGRM.

Now we show the approach to calculate the contributions of each branch to the eigenvectors of eGRM. Same as using *C* or 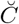, we first perform an eigenvector decomposition of the tree eGRM:

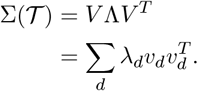

Then we can perform a linear regression:

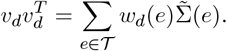

Since for large matrices there is no analytical solution for eigenvectors, it is impossible to mathematically derive *w*_*d*_(*e*) in a closed form. Therefore, in practice, the only way to calculate the contributions of each branch to the eigenvectors of a general eGRM would be to perform a linear regression without an intercept.

### 2 Relationship between eGRM and variance covariance matrix

From equations (1) to (3), we know that:

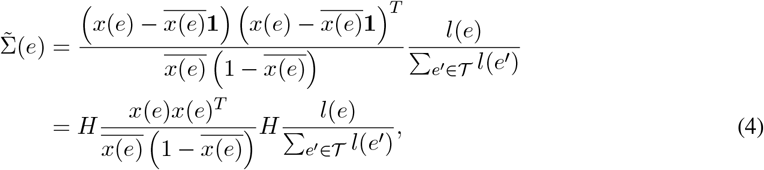

with

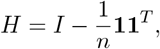

where *I* is the identity matrix and *n* is the number of tips of the tree 𝒯.

Traditionally, phylogeneticists use a variance-covariance matrix (VCV or *C*) by assuming that traits evolve following a Brownian motion across the tree. As variance-covariance matrix explicitly models the evolution of traits, it is natural to consider using variance-covariance matrix to compute eigenvectors for covariates in phylogenetic regression, which is shown in our previous paper [7]. Similar to eGRM, here we propose a branch version of a variance-covariance matrix:

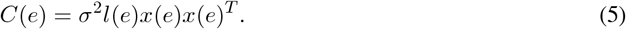

The intuition is that if tips *i* and *j* are both descendants of a branch *e*, then the branch would contribute *σ*^2^*l*(*e*) to their covariance under a Brownian motion, and thus we would have:

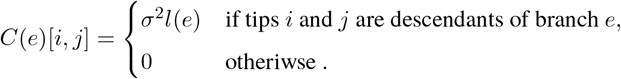

Note that this equation also works when *i* and *j* happen to be the same individual. The variance-covariance matrix for the whole tree 𝒯 would then be the sum of variance-covariance matrices for the branches:

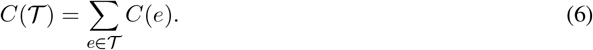

Combining equations (4) and (5), we can show:

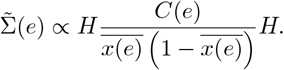

From the equation above, we observe that the differences between variance-covariance matrix and (weighted) eGRM of a branch are two-fold. First, the variance-covariance matrix does not mean-center the data, while eGRM is mean-centered both row- and column-wise. Second, variance-covariance matrix does not normalize the data whereas eGRM does.

We argue that mean-centering the matrix is an important step for including eigenvectors into phylogenetic regressions. In Principal Component Analysis (PCA), one would mean-center the data before calculating the variance-covariance matrix and performing the eigenvector decomposition; otherwise, the data could be biased by the mean. As eGRM is obtained after mean-centering the data *x*(*e*), performing an eigenvector decomposition on eGRM would be equivalent to performing PCA on the original data, except that the data are also normalized, such that the branches that give rise to more uneven partitions (*x*(*e*) being mostly zeroes or ones) among the tip species are given more weights. Since the variance-covariance matrix is obtained without mean-centering the data first, directly performing an eigenvector decomposition can generate biased PCs. Here we show how eigenvector decomposition can change before and after double-centering.

Suppose that the eigenvector decomposition of the variance-covariance matrix is:

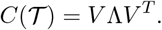

Then the doubled-centered variance-covariance matrix would be:

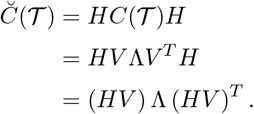

Note that this is not an eigenvector decomposition of 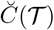, because the columns of *HV* are not orthonormal. Now the *i*-th column of the matrix would be:

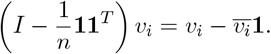

And thus we have:

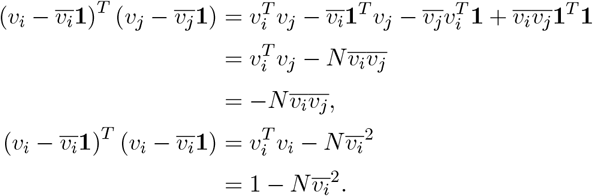

While the above shows that the eigenvector decomposition of 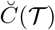 cannot be obtained via a simple transformation from the *C*(𝒯), there is actually no analytical relationship between the eigenvector decomposition of 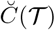 and that of *C*(T).

McVean [6] has suggested using *E*(*M*) as the matrix for eigenvector decomposition, with:

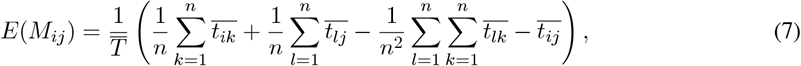

where 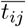 is the expected coalescence time for tips *i* and *j*, and 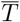 is the expected total branch length in the tree. Note that McVean’s *E*(*M*) is derived from expected coalescence times, where the tree is not fixed. However, assuming that the tree is fixed, we eventually obtain the same matrix as the double-centered variance-covariance matrix. We have:

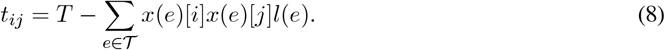

The intuition is that the distance from the most recent common ancestor of tips *i* and *j* to the root of the tree would be the sum of the lengths of branches that have both *i* and *j* as descendants.

Plugging equation (8) into (7), we get:

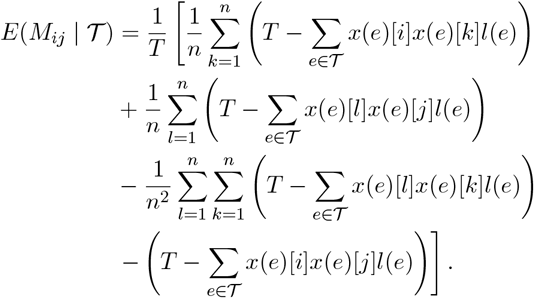

Simplifying the above equation gives us:

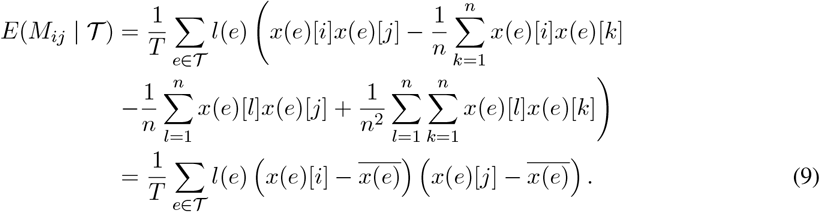

We can rewrite equation (9) in matrix format:

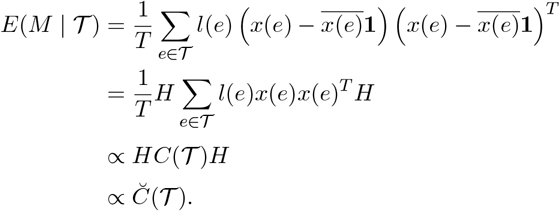

Therefore, we have shown that McVean’s *E*(*M*) is proportional to 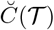 if the tree 𝒯 is fixed, and thus performing an eigenvector decomposition of *E*(*M* | 𝒯) would be equivalent to performing the same analyses on 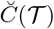.

### 3 Relationship between eigenvectors of distance matrix and variance-covariance matrix

When the method of using eigenvectors in phylogenetic regressions (PVR) was first proposed, Diniz-Filho et al. [2] uses a double-centered distance matrix to get the eigenvectors. Here we show that this is equivalent to using a double-centered variance-covariance matrix.

The phylogenetic distance between tips *i* and *j* would be:

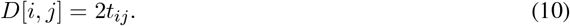

We can also rewrite equation (8) in a matrix format:

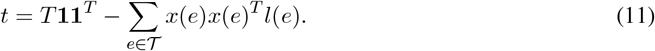

Combining equations (10) and (11), and assuming a Brownian motion *σ*^2^ of 1, we can derive the formula for the phylogenetic distance matrix of tree 𝒯:

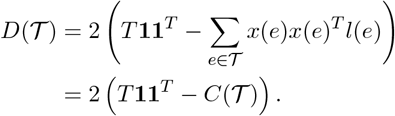

Now we show that if a vector *v* is an eigenvector of double-centered variance-covariance matrix 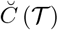, then it is also an eigenvector of double-centered distance matrix 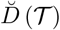.

The fact that a vector *v* is an eigenvector of 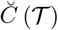 means there exists some eigenvalue *λ*_1_ such that:

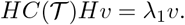

We need to show that there exists some eigenvalue *λ*_2_ such that:

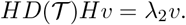

That is:

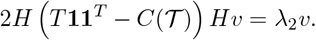

We only need to show that there exists some eigenvalue *λ*_3_ such that:

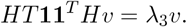

We note that the matrix on the left is a zero matrix because:s

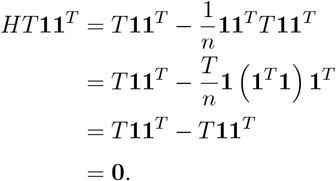

Therefore, *v* is an eigenvector for *HT* **11**^*T*^ *H* with corresponding eigenvalue *λ*_3_ = 0, and thus *v* is also an eigenvector for 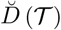 with corresponding eigenvalue *λ*_2_ = −2*λ*_1_. One thing to note here is that 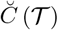 and 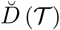 would always differ in the signs of their eigenvalues. In fact, we can show that 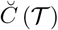 would always generate non-negative eigenvalues, whereas 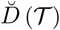 would always give non-negative eigenvalues.

To prove this, we can simply show that for any vector *v*:

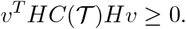

Using the definition of variance-covariance matrix in equations (5) and (6), we have:

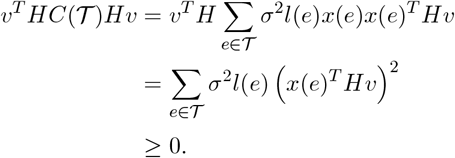

### 4 Eigenvectors of matrices for non-ultrametric trees

In non-ultrametric trees, the variance-covariance matrix *C* of the trait for tree 𝒯 can be computed following exactly the same equations (5) and (6), and after double-centering the variance-covariance matrix, the eigenvalues would still be all non-negative. However, when it comes to the distance matrix, since tips have different distances from the root of the tree, the equations in Section 3 cannot be applied. Here, we extend the discussion in Section 3 to non-ultrametric trees and show that after double-centering, the distance matrix and the variance-covariance matrix of the same tree still share the same eigenvectors.

First, we derive the distance matrix *D* of a tree 𝒯. de Vienne et al. [1] have shown that the phylogenetic distance between two tips *i* and *j* of a tree can be expressed using the heights of the nodes (distances from nodes to the root of the tree). Let *h*_*i*_, *h*_*j*_, and *h*_*MRCA*[*i,j*]_ be the heights of *i, j*, and their most recent common anestor, respectively. Then we have:

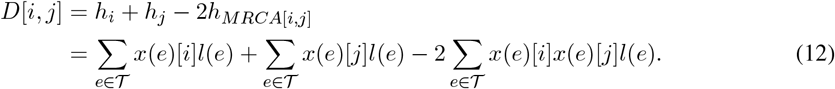

We can rewrite equation (12) in a matrix format:

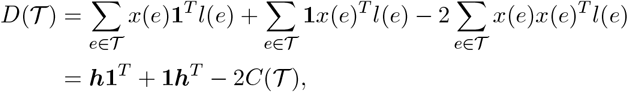

where the last line is obtained by assuming a variance of 1 for the Brownian motion and using ***h*** = ∑ _*e*∈ *𝒯*_ *x*(*e*)*l*(*e*) to denote the vector for heights of the tips.

Similar to Section 3, we can show that if a vector *v* is an eigenvector of 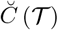, then it is also an eigenvector of 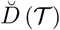. We only need to show that *v* is an eigenvector of double-centered matrix *H* ***h*1**^*T*^ + **1*h*** *H*. We note that:

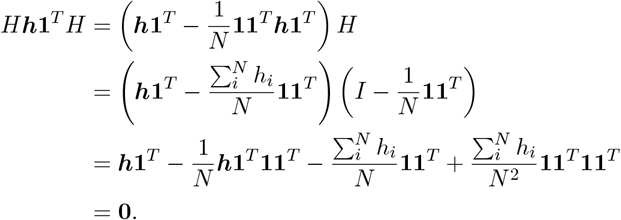

We also have:

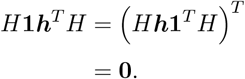

The calculations above give us:

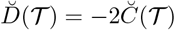

Therefore, we can show that *v* is an eigenvector of double-centered matrix *H* (*h***1**^*T*^ + **1***h*^*T*^) *H* with eigenvalue 0, and thus *v* is also an eigenvector of 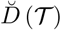 with eigenvalue −2*λ*_1_, where *λ*_1_ is the corresponding eigenvalue for *v* for 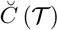.

de Vienne et al. [1] have pointed out that for non-ultrametric phylogenetic trees, the distance matrix *D* can be non-Euclidean, and they suggest using matrix *D*^∗^, which is obtained by taking the element-wise square root of *D*. Since this is a non-linear transformation, we cannot mathematically show the relationship between eigenvectors of double-centered distance matrix 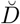 and double-centered element-wise square root distance matrix 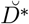. However, by the definition of a Euclidean distance matrix *D*^∗^, one can represent the *n* tips in R^*n*^ space such that:

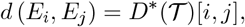

where *E*_*i*_ and *E*_*j*_ are the embeddings of tips *i* and *j*. To compute the embeddings matrix *E* from the distance matrix *D*^∗^, we first square the matrix element-wise and then perform the double-centering transformation and multiply the matrix by a constant of 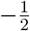. This gives us:

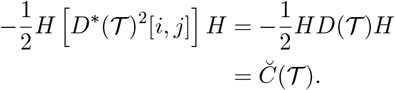

Next, we perform eigenvector-decomposition of the double-centered variance-covariance matrix and the embeddings of the *n* tips can be expressed using the eigenvectors and eigenvalues:

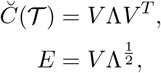

where *E* represents the embeddings of the *n* tips, with *E*[*i, d*] representing the embedding of the *i*-th tip in the *d*-th dimension.

With the embeddings of *n* tips available, we can perform a PCA on the embeddings and use the projections onto the PC-axes to capture the phylogenetic structure. We note that the embeddings matrix *E* is already zero-centered dimension-wise (column-wise). That is,

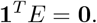

This is because:

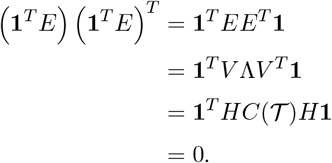

The covariance matrix for PCA is:

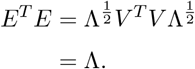

Therefore, the eigenvalues of *E*^*T*^ *E* are just the diagonal elements of Λ (the eigenvalues of the double-centered variance-covariance matrix), and the eigenvectors are just the standard basis vectors {*e*_*i*_ ∈ ℝ^*n*^}.

Now we show that using the projections of the embeddings of the tips onto the PC-axes is equivalent to using the eigenvectors of the double-centered variance-covariance matrix. Following what we have shown above, the projection of the embeddings of the tips onto the *d*-th PC-axis is *Ee*_*d*_. Therefore, we only need to show that *Ee*_*d*_ is the *d*-th leading eigenvector of doubled-centered variance-covariance matrix:

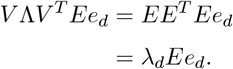

### 5 Notes on using non-centered variance-covariance matrix

Above we have shown the correctness of using eigenvectors of a double-centered variance-covariance matrix 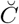 to correct for confounding in phylogenetic regression. However, in practice, we can show that even though using a non-centered variance-covariance matrix *C* would generate different eigenvectors, in general it can lead to similar results.

As we have discussed in both Section 2 and 4, the double-centered 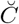 matrix for a tree with *n* tips is equivalent to the variance-covariance matrix of *n* points in ℝ^*n*^ space (relative to the mean embedding). The non-centered *C* then would be the variance-covariance matrix of *n* points in ℝ^*n*^ space relative to the original point **0**. Therefore, getting the eigenvectors of the non-centered *C* would have the same effect as not-centering the data when performing PCA. In PCA, if one does not perform mean-centering on the data, a large part of the variance from the origin would come from the mean vector, so the first few eigenvectors would be strongly biased by the mean of the data, and the eigenvalues would be overestimated.

However, eigenvectors corresponding to smaller eigenvalues are in general orthogonal to the mean vector, and are thus less affected by the mean-centering transformation. For these smaller eigenvalues, we observe that in general the *i*-th eigenvalue and eigenvector pair of a double-centered 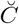 is similar to the (*i*+1)-th eigenvalue and eigenvector pair of a non-centered *C*. The last eigenvalue and eigenvector pair (0 and **1**) of the doubled-centered 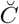 does not have a corresponding pair in the non-centered *C*, because an eigenvalue of 0 means that the last eigenvector does not explain any variance among the data.

Even though we cannot verify the observation above mathematically, we can think about this in terms of matrix perturbation theory. The difference between the double-centered 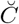 and the non-centered *C* is:

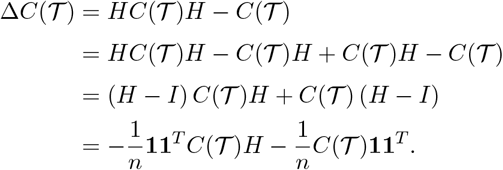

Note that **11**^*T*^ is a rank-1 matrix, so the two components involving **11**^*T*^ in the equation above also give rise to matrices of rank 1, and thus Δ*C*(𝒯) is a matrix of rank of at most 2, and at least *n* − 2 of its eigenvalues are 0. In fact, we can even show that the matrix has either rank 2 with one positive and one negative eigenvalue or rank 1 with one negative eigenvalue. This is equivalent to showing that matrix *M* = **11**^*T*^ *C*(𝒯)*H* + *C*(𝒯)**11**^*T*^ has either rank 2 with one positive and one negative eigenvalue or rank 1 with one positive eigenvalue.

First, we can show that *M* always has one positive eigenvalue using Rayleigh quotient. Let ***x*** = **1**. Then we have:

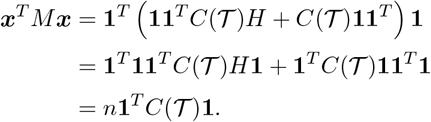

The Rayleigh quotient would be:

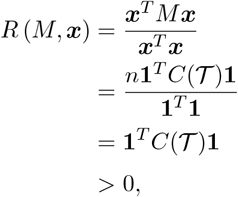

where we have the last line because *C*(𝒯) has non-negative entries and is not all zeroes. We know that:

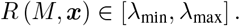

Therefore, we know that matrix *M* has at least one positive eigenvalue. Now we show that matrix *M* has at least one negative eigenvalue. We note that:

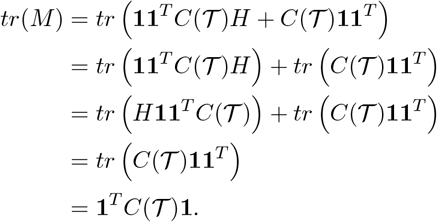

We also have:

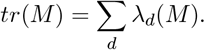

Then, we have:

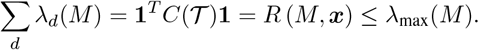

The equality above holds if and only if *M* has rank 1. Otherwise, *M* must have at least one negative eigenvalue such that the sum of all the eigenvalues add up to a value less than its largest eigenvalue. And since *M* has at most a rank of 2, in this case, *M* would have exactly one positive and one non-positive eigenvalue. Therefore, if we order the eigenvalues *λ*_1_, *λ*_2_, …, *λ*_*n*_ of Δ*C*(𝒯) in decreasing order, we obtain:

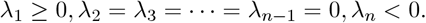

By Weyl’s Inequalities [8], we have:

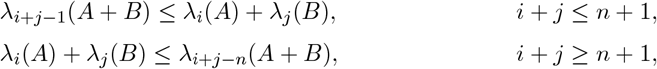

where *λ*_*i*_(*A*) is the *i*-th largest eigenvalue of matrix *A*. Let *A* be the non-centered *C*(𝒯) and *B* be Δ*C*(𝒯). Since *λ*_*j*_ (Δ*C*(𝒯)) is 0 for all *j* = 2, 3, …, *n* − 1, we choose *j* = 2 for the first inequality and *j* = *n* − 1 for the second. Then we can show that:

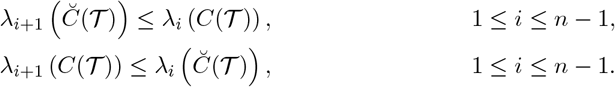

This sandwich pattern of the eigenvalues of *C* and 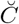 indicates that for later dimensions, where the eigenvalues are typically small, not double-centering the matrix *C* would not impact the eigenvalues by much.

The last piece of the puzzle comes from Equation (18) from our previous paper [7], where we have:

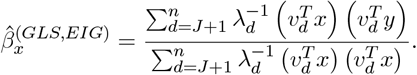

Therefore, even when the eigenvectors of a non-centered variance-covariance matrix are used in phylogenetic regressions, when getting rid of the first *J* eigenvectors, one also gets rid of the eigenvectors that are the most biased by mean of the matrix. Since the remaining eigenvalue and eigenvector pairs of the non-centered variance-covariance matrix are similar to the one of the double-centered variance-covariance matrix, the estimate of *β* in the analyses would not deviate much from the estimate using a double-centered variance-covariance matrix.

### 6 Overall contributions of each branch

Once the contributions of the branches to each of the eigenvectors are obtained, we can also compute the overall contributions of each branch. Recall that *w*_*d*_(*e*) is the contribution of some branch *e* of a phylogenetic tree to the *d*-th eigenvector of some version of variance-covariance matrix of the tree. Then the overall contribution of the branch *e* across all the dimensions would be:

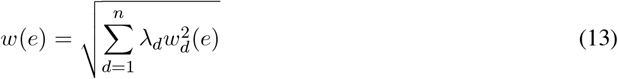

where *λ*_*d*_ is the *d*-th eigenvalue of the variance-covariance matrix. Note that the formula looks very much like a Euclidean norm, except that each dimension is also weighted by the corresponding eigenvalue. We choose the Euclidean norm for two reasons: (1) the eigenvectors represent dimensions orthogonal to each other, which makes the Euclidean norm appropriate, (2) *w*_*d*_(*e*) can be negative, so directly adding up the contributions across the dimensions could potentially cancel out some of the terms and underestimate the contributions of some branches. We also weight each dimension by *λ*_*d*_ because the first few dimensions explain the most variance and are thus more important than the later dimensions. The overall contributions of each branch can be used as a metric for the importance of each branch in the phylogenetic tree.

### 7 Connection between PGLS and PGLMM

We use PGLS and PGLMM interchangeably throughout the discussion in our paper, because they are equivalent in our simulation scenarios. Here we provide the proof. Suppose that we are including the first *J* eigenvectors as covariates, then in PGLS, we have:

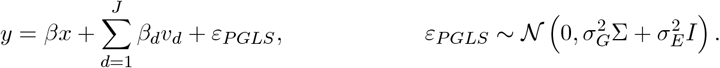

In PGLMM, we have:

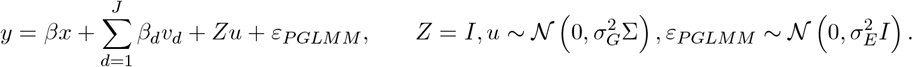

The reason why the design matrix *Z* above is equivalent to the identity matrix is that we have only one individual for each species. Then, for both equations above, we can show that *y* follows the same multivariate normal distribution:

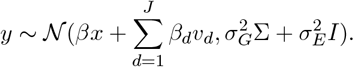

The only difference between the two methods is that PGLS is including the variance-covariance matrix Σ in the variance structure of the residuals, whereas PGLMM is including Σ as random effects.

### 8 Performance of different methods in Felsenstein’s Worst Case

In our simulations for Felsenstein’s Worst Case (FWC), we observe that when we include *v*_1_ of *C* as fixed effect, the results for OLS and PGLS are exactly the same. Here we show mathematically why this would be the case. Suppose that we have *n* species in each clade in FWC, and we assume a Brownian motion rate of 1, then the phylogenetic variance-covariance matrix for the 2*n* species would be:

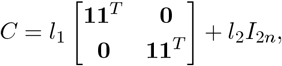

where **1** is a one-vector of length *n*, **0** is an *n* × *n* zero-matrix, *I*_2*n*_ is the 2*n* × 2*n* identity matrix, and *l*_1_ and *l*_2_ are the lengths of the branches outside and within the cade, respectively.

Note that we can rewrite *C* as:

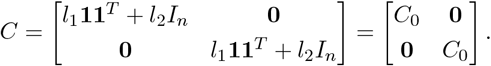

Now we show that if *λ* and *v* is an eigenvalue-eigenvector pair for *C*_0_, then *λ* and [*v*^*T*^ 0 0 … 0]^*T*^ and *λ* and [0 0 … 0 *v*^*T*^]^*T*^ are two pairs of eigenvalues and eigenvectors for *C*. This is because:

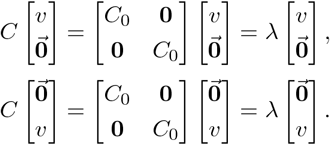

Thus, to get the eigenvector decomposition for *C*, we only need to do the decomposition on *C*_0_. We can easily show that *l*_1_*n* + *l*_2_ and **1** is a pair of eigenvalue and eigenvector for *C*_0_, since:

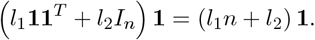

Similarly, we can show that *l*_2_ and any vector *v* such that *v*^*T*^ **1** = 0 is a pair of eigenvalue and eigenvector for *C*_0_, since:

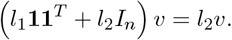

There would be a total of *n*−1 such eigenvectors *v* in ℝ^*n*^ space. Thus, the eigenvalues of *C*_0_ are *l*_1_*n*+*l*_2_, *l*_2_, *l*_2_, …, *l*_2_, and the eigenvalues of *C* are just the duplicates of these eigenvalues. Now, for OLS with *v*_1_ as covariate, our design matrix would be *X* = **1** *x v*_1_, which is slightly different from the formula used to derive equation (16) in our previous paper [7]. However, we observe that *y* = *β*_0_ + *β*_*x*_*x* + *β*_1_*v*_1_ + *ε*_*OLS*_ is equivalent to *y* = *β*_*x*_*x* + *β*_1_*v*_1_ + *β*_2_*v*_2_ + *ε*_*OLS*_, since:

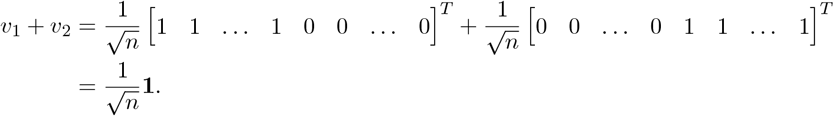

That is, including the first eigenvector and the intercept as fixed effects in FWC is the same as including the first two eigenvectors without the intercept. And thus we have:

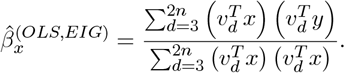

Similarly, we can plug in equation (18) in our previous paper [7] using the same trick for PGLS:

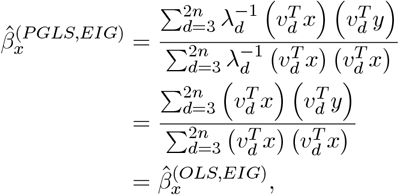

where the second row is obtained by plugging in *v*_*d*_ = *l*_2_ for all *d* ≥ 3. Therefore, we have shown using mathematical analyses why we would obtain exactly the same results for OLS and PGLS when including the first eigenvector and the intercept as covariates.

### 9 Supplementary Methods

In the sections above, we have addressed multiple matrices that can be used to capture the relationships between species of a phylogenetic tree, so we first compared how branches contribute to the eigenvectors of these different matrices, including expected genetic relatedness matrix (eGRM, or Σ), phylogenetic variance-covariance matrix (VCV, or *C*), and its double-centered version (dcVCV, or 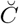). We specifically investigated three different types of trees. For Felsenstein’s Worst Case (FWC), we checked how the branches contribute to *v*_1_ of these different matrices (Fig. S1). We then looked at the contributions of branches to the first three eigenvectors of these different matrices for a 10-species Yule tree (Fig. S2) and a 10-species caterpillar tree (Fig. S3).

While we have shown in the sections above that it is theoretically rigorous to use the double-centered version 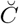 to get the eigenvectors, we also want to know how using a non-centered *C* in practice might change the results of phylogenetic regressions. To compare the performance of using *C* and 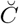 to get the eigenvectors, we simulated Yule trees, caterpillar trees, fully balanced trees, and coalescent trees of 128 species. For each tree type, we generated 500 different ultrametric trees with varying branch lengths (Fig. S4). Then we simulated *x* and *y* as arising independently from a Brownian motion along each of the trees. We compared the performance of using different numbers of eigenvectors of *C* and 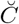 in OLS and PGLS for phylogenetic regression with `R` function `lm()` and `phylolm()` from the package `phylolm` [4, 5].

We also checked the performance of using the eigenvectors of a non-centered *C* in phylogenetic regressions for non-ultrametric trees, where we followed similar procedure as in the ultrametric case, but we used pure caterpillar trees, fully balanced trees, and moderately imbalanced trees of 128 species instead (Fig. S5). The moderately imbalanced trees were simulated by first generating coalescent trees and then reassigning branch lengths as independent exponential random variables Exp(1).

Similar to Fig. 2, we tested the effect of tree size on the performance of OLS and PGLS with eigenvectors in non-ultrametric trees. We simulated caterpillar trees, fully balanced trees, and moderately imbalanced trees of 8, 16, 32, 64, 128, 256, 512, and 1024 species (Fig. S6). For each combination of tree type and tree size, we generated 500 different non-ultrametric trees with varying branch lengths. The rest of the procedure is exactly the same as in previous simulations.

To understand the number of eigenvectors we need to include as covariates in the model to control for confounding when there are non-Brownian shifts on a phylogenetic tree, we generated the same 16-species Yule tree as in Fig. 4, and we visualized the contributions of branches to the first eight eigenvectors of the phylogenetic variance-covariance matrix *C* (Fig. S7). We first looked at cases when there is only one non-Brownian shift on the tree. The simulation procedure is exactly the same as is mentioned in the Methods section for the main text. For each combination of shift position, shift type, and shift size, we ran a total of 100 replicates (Figs. S8 and S9).

We further investigated how the performance of PGLS with no eigenvectors can be impacted by where the shift falls on the tree. We generated a total of 500 Yule trees of 16 species. For each tree, we first calculated the overall contributions of the branches across the dimensions using equation (13) from Section 6, with which we ranked the branches in decreasing order. Then we simulated *x* and *y* following the same procedure, and for each combination of shift type and size, we checked how the false positive rate of PGLS changes with the rank of the branch where the shift occurs (Fig. S10).

Next we turned to investigate the cases when there are two non-Brownian shifts on the tree. We used the same 16-species Yule tree as in the cases for one non-Brownian shift in Fig. S7. Similar to previous simulations, we first simulated *x* and *y* as arising independently from a Brownian motion along each of the trees. Then, out of the nine branches that contribute mostly to the first eight eigenvectors of the *C* matrix (branches labeled 1, 2, 11, 15, 17, 23, 24, 25, and 26 in Fig. S7), we selected two branches when the shifts occur. Then we evaluated the performance of using different numbers of eigenvectors of *C* in PGLS. For each combination of shift position pairs, shift type, and shift size, we ran a total of 100 replicates (Figs. S11 and S12).

We also analyzed the cases when there are multiple non-Brownian shifts on the same 16-species Yule tree as in Fig. S7. We specifically looked at the cases when there are shifts on 4 branches (labeled 1, 2, 23, and 25 in Fig. 7), when there are shifts on 8 branches (labeled as 1, 2, 11, 15, 17, 23, 25, and 26), and when there shifts on 16 branches (labeled as 1, 2, 9, 10, 11, 15, 17, 18, 23, 24, 25, and 26). We ran the analyses using the same simulation procedure as in the cases for two shifts, and we ran 100 replicates for each combination of shift positions, shift type, and shift size (Fig. S13).

While we focused mostly on how the combined method helps mitigate confounding to reduce false positives in phylogenetic regressions in all the previous simulations, we are also interested in the performance of the combined method in terms of power. We first compared the performance of using the eigenvectors of *C* and 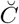 in OLS and PGLS. We simulated Yule trees, caterpillar trees, fully balanced trees, and coalescent trees of 128 species. For each tree type, we generated 500 different ultrametric trees with varying branch lengths. Then we simulated *x* as arising from a Brownian motion along each of the trees, and *y* as the combination of a dependent part *βx* and an independent part *y*_0_, where *β*, the true regression coefficient, was set to be 0.5, and *y*_0_ was simulated as arising from a Brownian motion along the tree independent from *x*. Then we compared the recall (1 − False Negative Rate) of using different numbers of eigenvectors of *C* and 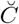 in OLS and PGLS for phylogenetic regressions (Fig. S14).

Next we explored the effect of tree size and *β* on the recall of using different numbers of eigenvectors in OLS and PGLS. We simulated Yule trees of 8, 16, 32, 64, 128, 256, 512, and 1024 species (Fig. S15). Then we simulated *x* as arising from a Brownian motion along each of the trees, and *y* as the combination of a dependent part *βx* and an independent part *y*_0_, where *β*, the true regression coefficient, was set to be some value from 0.0625 to 16, and *y*_0_ was simulated as arising from a Brownian motion along the tree independent from *x*. Then we compared the recall of using different numbers of eigenvectors of *C* in OLS and PGLS. For each combination of tree size and *β*, we ran a total of 100 different replicates.

Finally, we looked at the recall when there is also a non-Brownian shift on the tree, and we used the same 16-species Yule tree as in Fig. S7. We focused on a subset of branches where the shifts may occur (labeled as 1, 2, 23, and 25 in Fig. S7). We simulated *x* as arising from a Brownian motion along the tree and generated the shift by adding a normal random variable (or a constant) to the species that are descendants of the branch where the shift occurs. Then we simulated *y* as the combination of a dependent part *βx* and an independent part *y*_0_, where *β*, the true regression coefficient, was set to be some value from 0.0625 to 2, and *y*_0_ was simulated as arising from a Brownian motion along the tree plus some shift, both of which are independent from *x*. Next we compared the recall of using different numbers of eigenvectors of *C* in OLS and PGLS, and for each combination of shift position, shift type, shift size, and *β*, we ran a total of 100 different replicates (Figs. S16 and S17).

## Supplementary Figures

**Fig S1:**
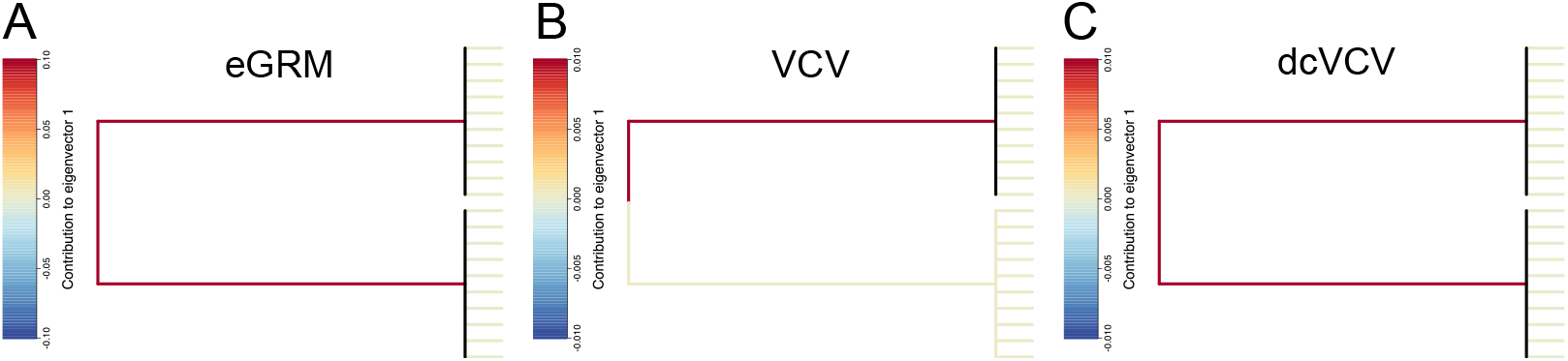
Contributions of branches to the first eigenvector *v*_1_ of different matrices of a phylogenetic tree depicting Felsenstein’s Worst Case (FWC). Each clade contains 10 species. (A) Contributions of branches to *v*_1_ of the expected genetic relatedness matrix (Σ) of the phylogenetic tree depicting FWC. (B) Contributions of branches to *v*_1_ of the non-centered phylogenetic variance-covariance matrix (*C*) of the phylogenetic tree depicting FWC. (C) Contributions of branches to *v*_1_ of the double-centered phylogenetic variance-covariance matrix 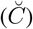 of the phylogenetic tree depicting FWC. The vertical branches are black because the tip branches slightly differ in the their contributions to *v*_1_.

**Fig S2:**
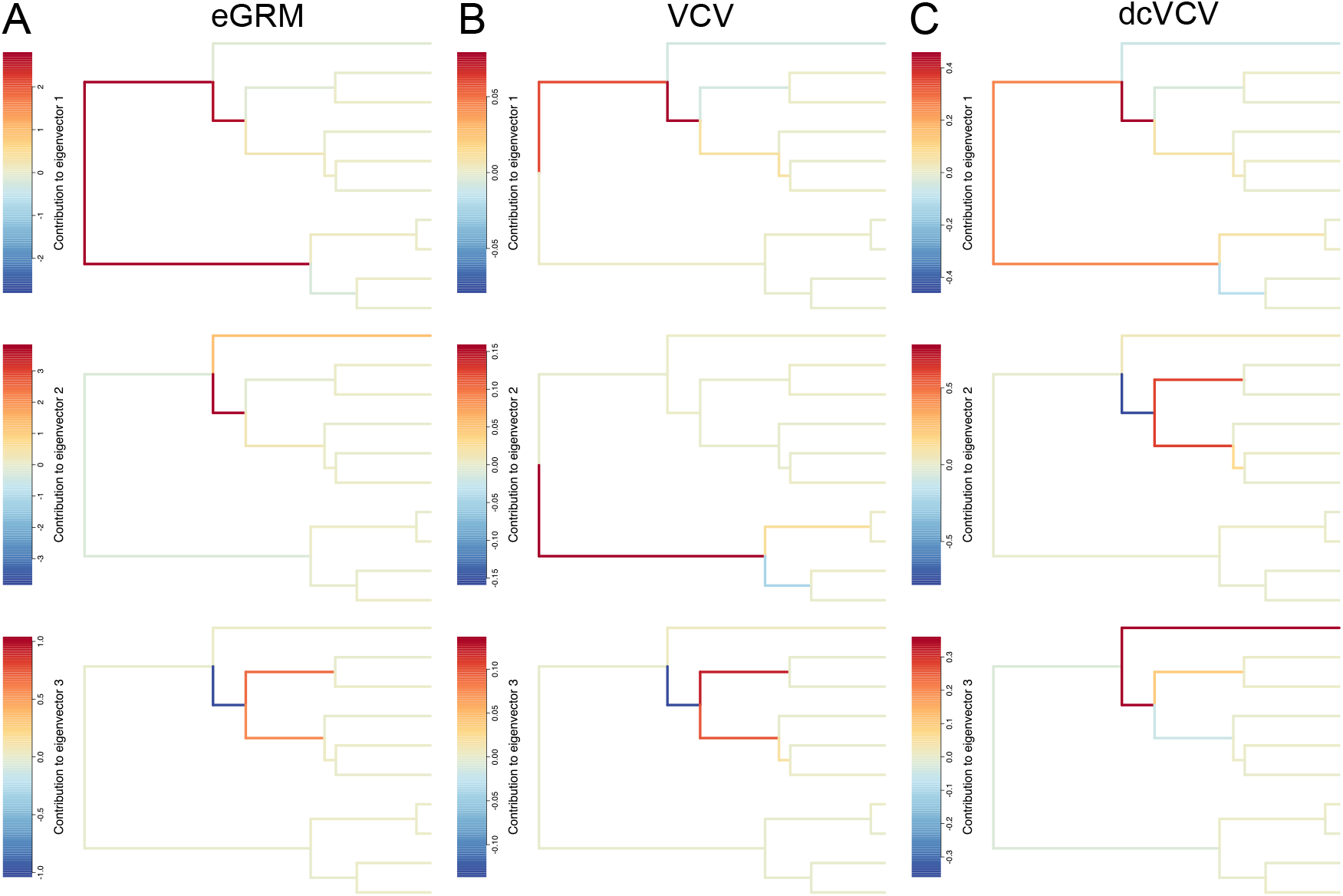
Contributions of branches to the first three eigenvectors of different matrices of a Yule tree containing 10 species. (A) Contributions of branches to the first three eigenvectors of the expected genetic relatedness matrix (Σ) of the Yule tree. (B) Contributions of branches to the first three eigenvectors of the non-centered phylogenetic variance-covariance matrix (*C*) of the Yule tree. (C) Contributions of branches to the first three eigenvectors of the double-centered phylogenetic variance-covariance matrix 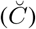 of the Yule tree.

**Fig S3:**
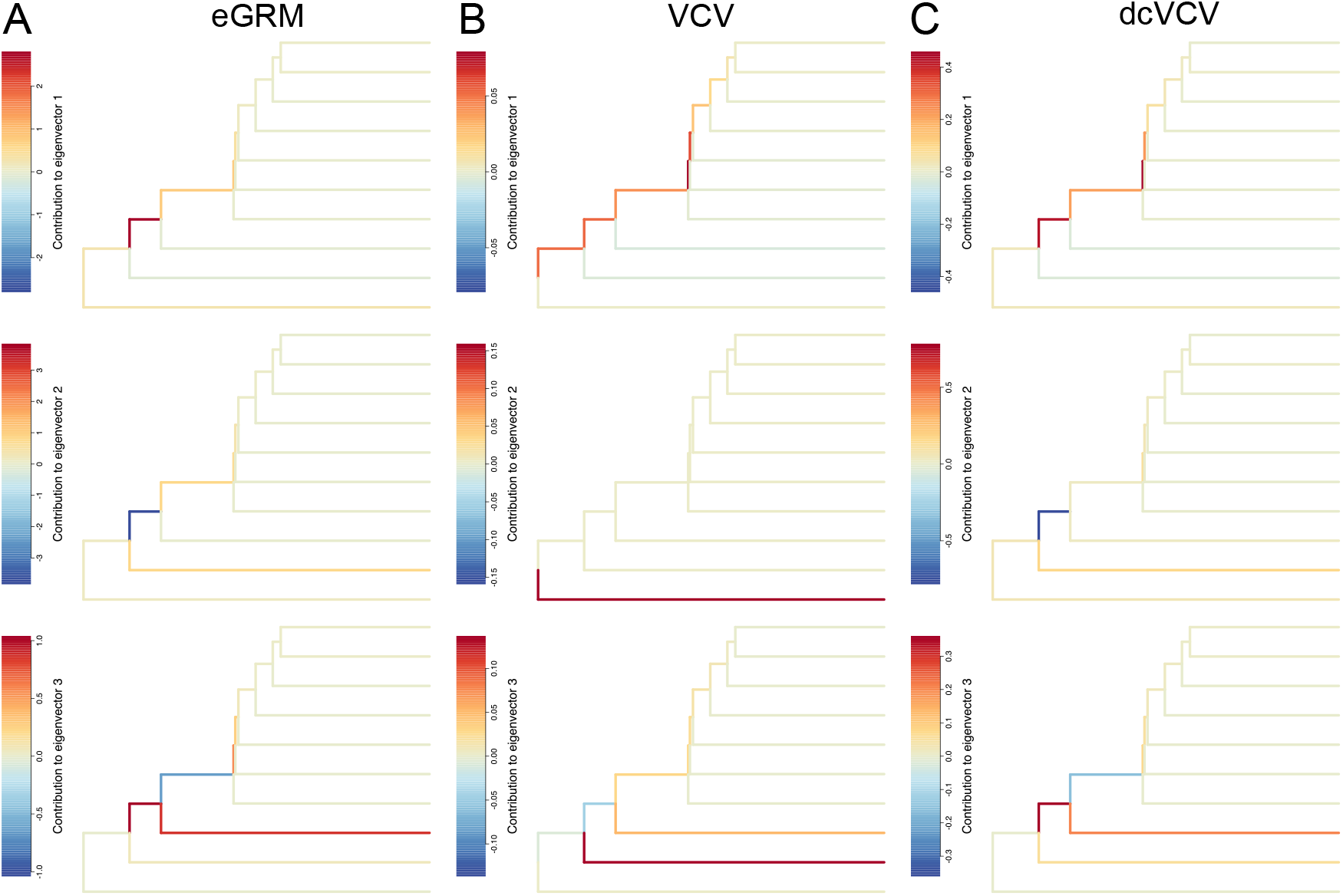
Contributions of branches to the first three eigenvectors of different matrices of a caterpillar tree containing 10 species. (A) Contributions of branches to the first three eigenvectors of the expected genetic relatedness matrix (Σ) of the caterpillar tree. (B) Contributions of branches to the first three eigenvectors of the non-centered phylogenetic variance-covariance matrix (*C*) of the caterpillar tree. (C) Contributions of branches to the first three eigenvectors of the double-centered phylogenetic variance-covariance matrix 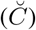 of the caterpillar tree.

**Fig S4:**
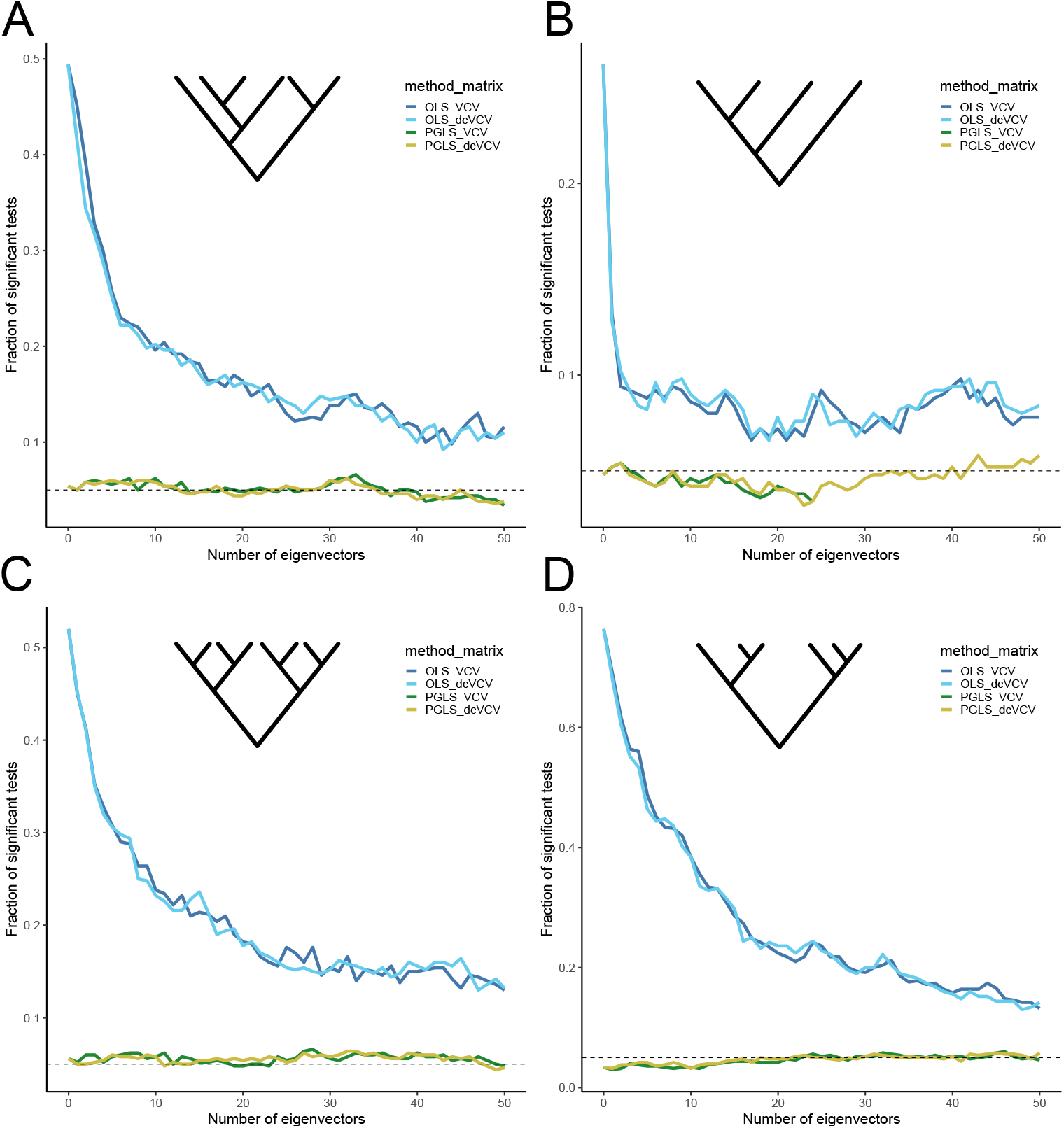
The performance of using eigenvectors of non-centered and double-centered phylogenetic variance-covariance matrices (*C* and 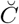) in ordinary least squares (OLS) and phylogenetic generalized least squares (PGLS) to remove phylogenetic confounding for different types of ultrametric trees. (A) Performance of OLS and PGLS with eigenvectors of *C* and 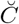 in models related by a Yule tree of 128 species. For simplicity, a Yule tree with only 6 tips is shown. The horizontal axis shows the number of eigenvectors included as covariates, and the vertical axis shows the fraction of tests that would be significant at the 0.05 level. (B) Performance of OLS and PGLS with eigenvectors of *C* and 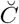 in models related by a caterpillar tree of 128 species. For simplicity, a caterpillar tree with only 4 tips is shown. The horizontal axis shows the number of eigenvectors included as covariates, and the vertical axis shows the fraction of tests that would be significant at the 0.05 level. (C) Performance of OLS and PGLS with eigenvectors of *C* and 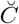 in models related by a fully balanced tree of 128 species. For simplicity, a fully balanced tree with only 8 tips is shown. The horizontal axis shows the number of eigenvectors included as covariates, and the vertical axis shows the fraction of tests that would be significant at the 0.05 level. (D) Performance of OLS and PGLS with eigenvectors of *C* and 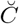 in models related by a coalescent tree of 128 species. For simplicity, a coalescent tree with only 6 tips is shown. The horizontal axis shows the number of eigenvectors included as covariates, and the vertical axis shows the fraction of tests that would be significant at the 0.05 level.

**Fig S5:**
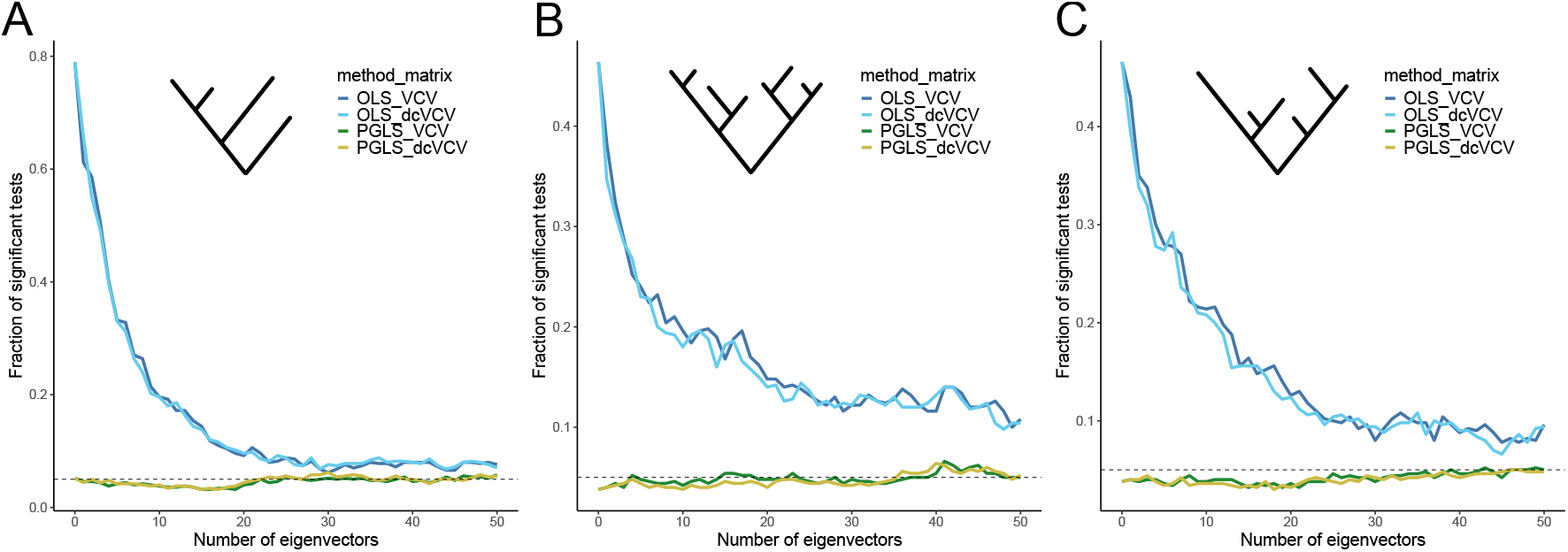
The performance of using eigenvectors of non-centered and double-centered phylogenetic variance-covariance matrices (*C* and 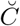) in ordinary least squares (OLS) and phylogenetic generalized least squares (PGLS) to remove phylogenetic confounding for different types of non-ultrametric trees. (A) Performance of OLS and PGLS with eigenvectors of *C* and 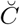 in models related by a non-ultrametric caterpillar tree of 128 species. For simplicity, a caterpillar tree with only 4 tips is shown. The horizontal axis shows the number of eigenvectors included as covariates, and the vertical axis shows the fraction of tests that would be significant at the 0.05 level. (B) Performance of OLS and PGLS with eigenvectors of *C* and 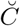 in models related by a non-ultrametric fully balanced tree of 128 species. For simplicity, a fully balanced tree with only 4 tips is shown. The horizontal axis shows the number of eigenvectors included as covariates, and the vertical axis shows the fraction of tests that would be significant at the 0.05 level. (C) Performance of OLS and PGLS with eigenvectors of *C* and 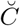 in models related by a non-ultrametric moderately imbalanced tree of 128 species. For simplicity, a moderately imbalanced tree with only 6 tips is shown. The horizontal axis shows the number of eigenvectors included as covariates, and the vertical axis shows the fraction of tests that would be significant at the 0.05 level.

**Fig S6:**
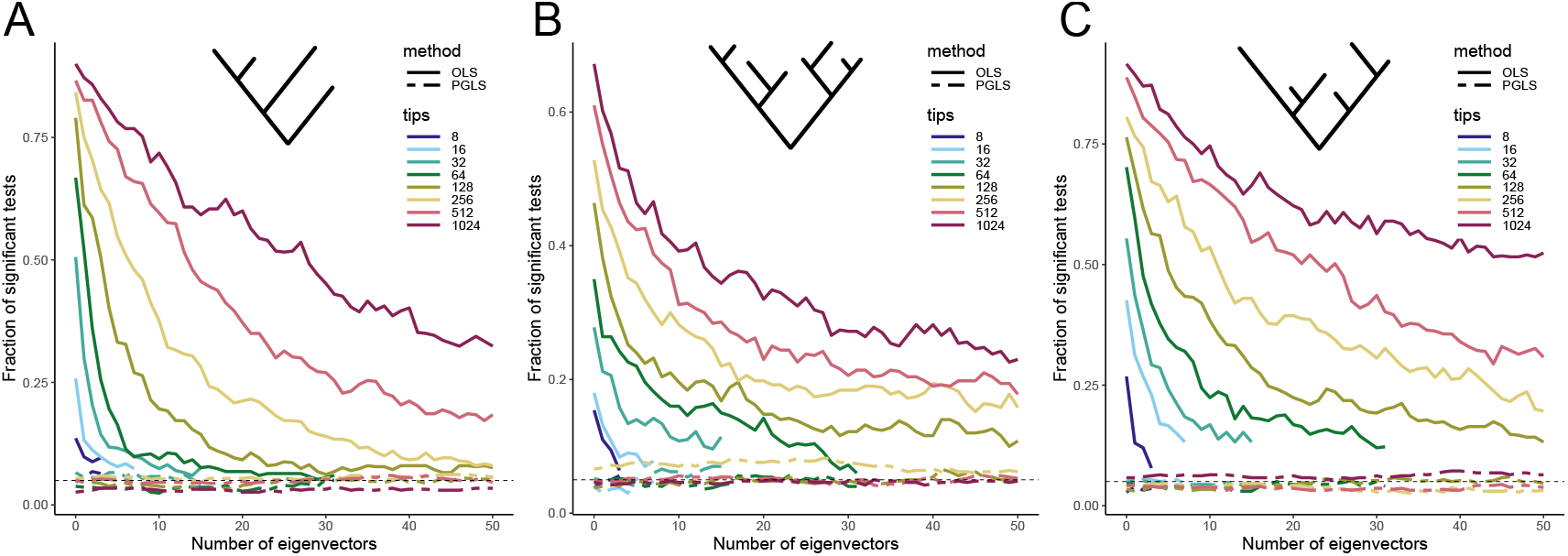
The false positive rate after adding eigenvectors in OLS and PGLS for different types of non-ultrametric trees when the data is generated by a pure Brownian motion with no additional shifts. (A) Performance of OLS and PGLS in models related by a non-ultrametric caterpillar tree of varying numbers of species. For simplicity, a non-ultrametric caterpillar tree with only 4 tips is shown. The horizontal axis shows the number of eigenvectors included as covariates, and the vertical axis shows the fraction of tests that would be significant at the 0.05 level. (B) Performance of OLS and PGLS in models related by a non-ultrametric fully balanced tree of varying numbers of species. For simplicity, a non-ultrametric fully balanced tree with only 8 tips is shown. The horizontal axis shows the number of eigenvectors included as covariates, and the vertical axis shows the fraction of tests that would be significant at the 0.05 level. (C) Performance of OLS and PGLS in models related by a non-ultrametric moderately imbalanced tree of varying numbers of species. For simplicity, a non-ultrametric moderately imbalanced tree with only 6 tips is shown. The horizontal axis shows the number of eigenvectors included as covariates, and the vertical axis shows the fraction of tests that would be significant at the 0.05 level.

**Fig S7:**
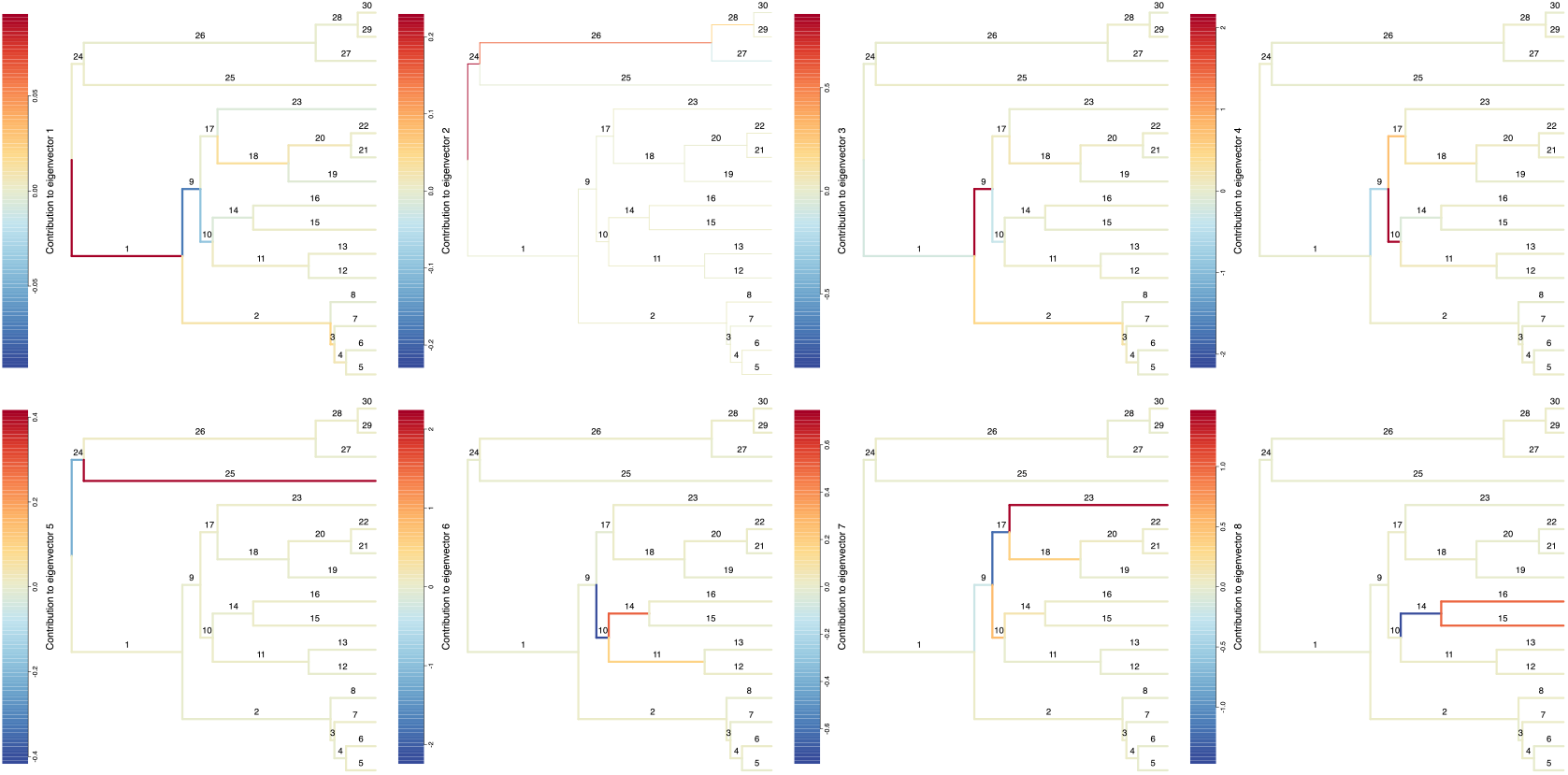
Contributions of branches to the first eight eigenvectors of the phylogenetic variance-covariance matrix *C* of a Yule tree containing 16 species. The numbers above each branch indicate the labels for the branches.

**Fig S8:**
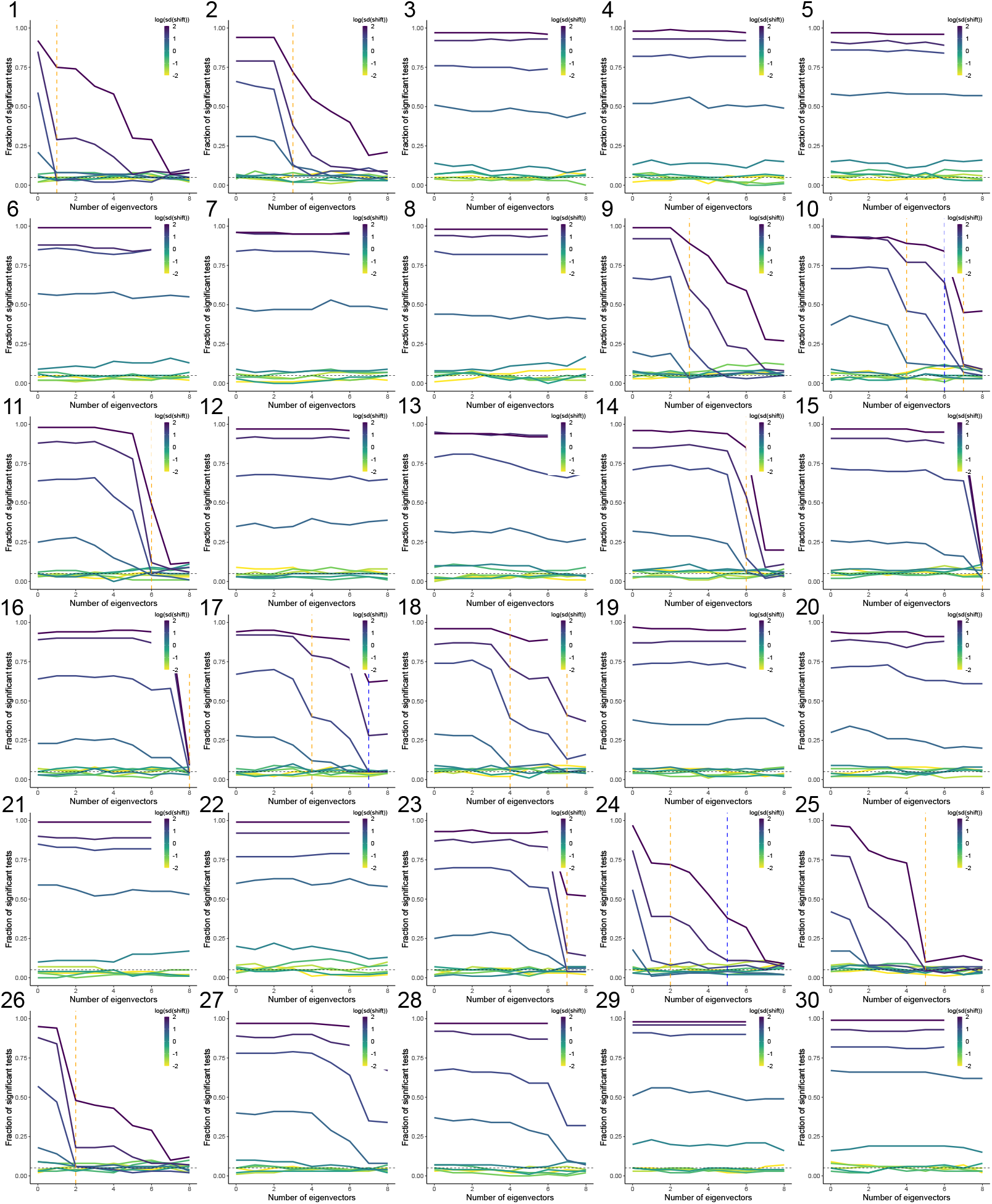
The performance of using eigenvectors in phylogenetic generalized least squares (PGLS) to remove phylogenetic confounding in a model with 16 tips related by a Yule tree with non-Brownian shifts in both predictor and outcome variables. The shift is simulated by adding a normal random variable with varying standard deviation to the species that are descendants of the branch. The label of each panel corresponds to the label of the branch where the shift occurs on the tree in Fig. S7. The horizontal axis shows the number of eigenvectors included as covariates, and the vertical axis shows the fraction of tests that would be significant at the 0.05 level. The vertical orange dashed lines show the dimensions to which the branch concerned makes large positive contributions, and the vertical blue dashed lines show the dimensions to which the branch concerned makes large negative contributions.

**Fig S9:**
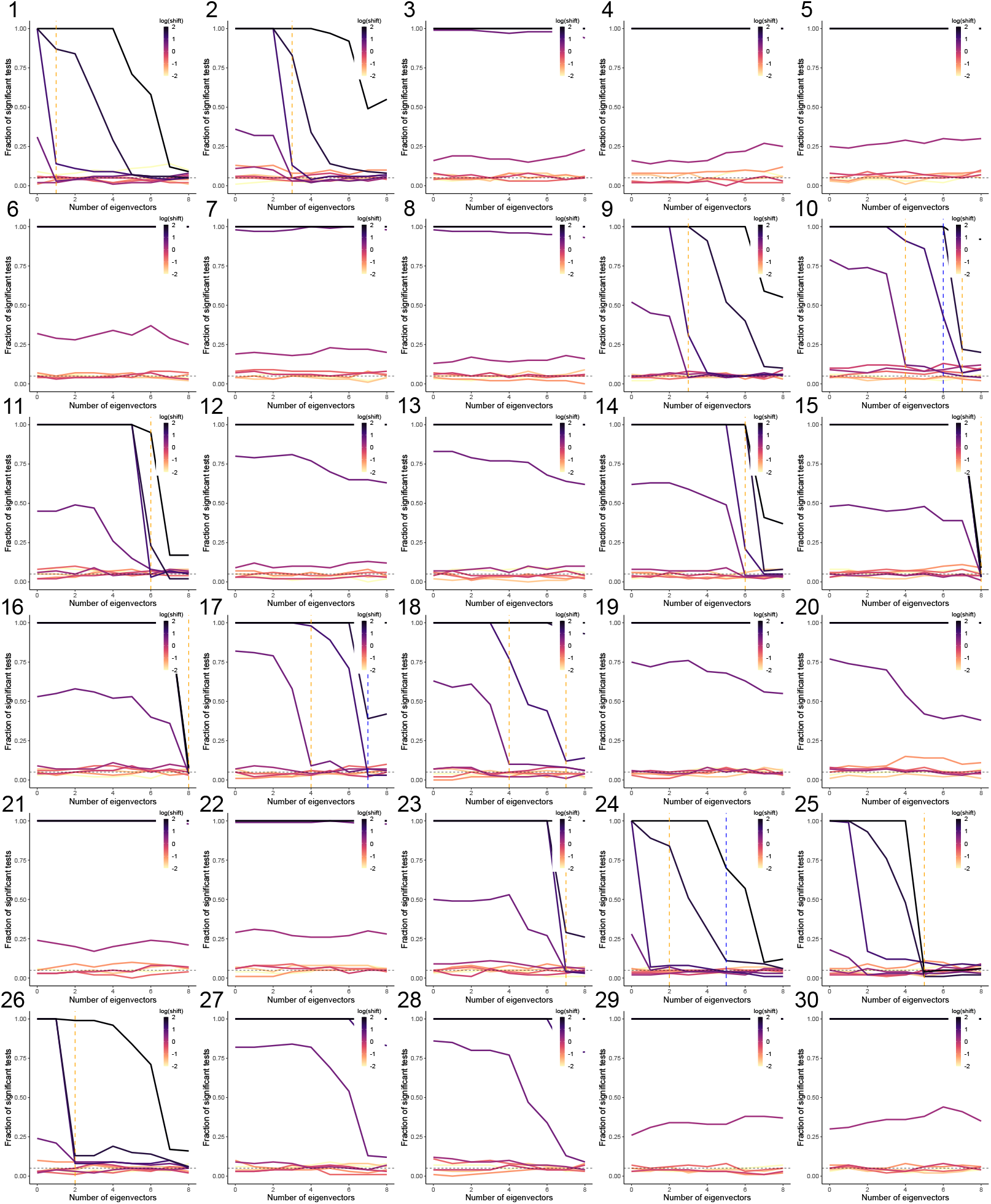
The performance of using eigenvectors in phylogenetic generalized least squares (PGLS) to remove phylogenetic confounding in a model with 16 tips related by a Yule tree with non-Brownian shifts in both predictor and outcome variables. The shift is simulated by adding a constant with varying size to the species that are descendants of the branch. The label of each panel corresponds to the label of the branch where the shift occurs on the tree in Fig. S7. The horizontal axis shows the number of eigenvectors included as covariates, and the vertical axis shows the fraction of tests that would be significant at the 0.05 level. The vertical orange dashed lines show the dimensions to which the branch concerned makes large positive contributions, and the vertical blue dashed lines show the dimensions to which the branch concerned makes large negative contributions.

**Fig S10:**
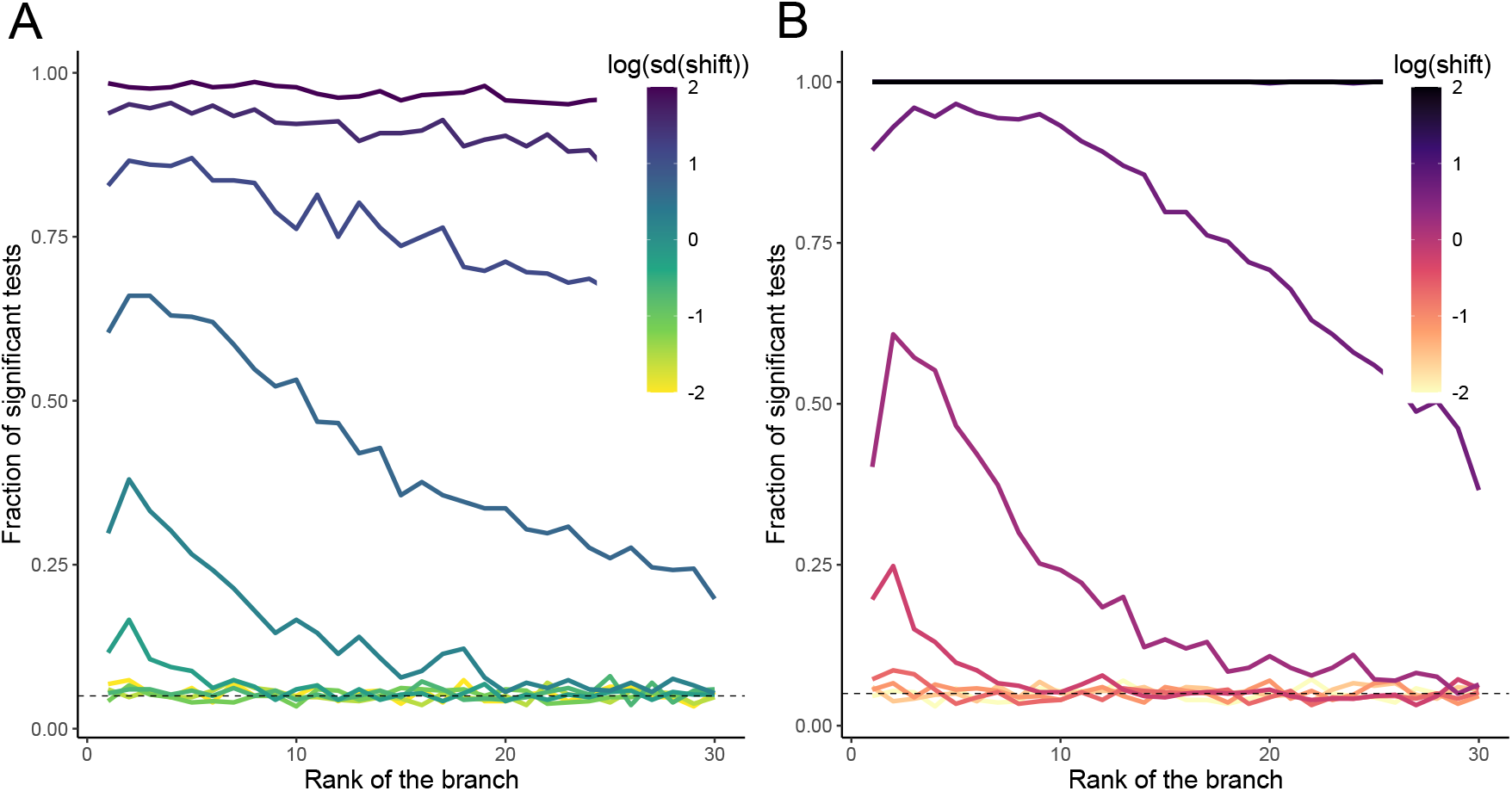
The performance of pure phylogenetic generalized least squares (PGLS) to remove phylogenetic confounding in models with 16 tips related by a Yule tree with a non-Brownian shift in both predictor and outcome variables. (A) The shift is simulated by adding a normal random variable with varying standard deviation to the species that are descendants of the branch. The horizontal axis shows the rank of the branches by overall contributions in decreasing order, and the vertical axis shows the fraction of tests that would be significant at the 0.05 level. (B) The shift is simulated by adding a constant with varying size to the species that are descendants of the branch. The horizontal axis shows the rank of the branches by overall contributions in decreasing order, and the vertical axis shows the fraction of tests that would be significant at the 0.05 level.

**Fig S11:**
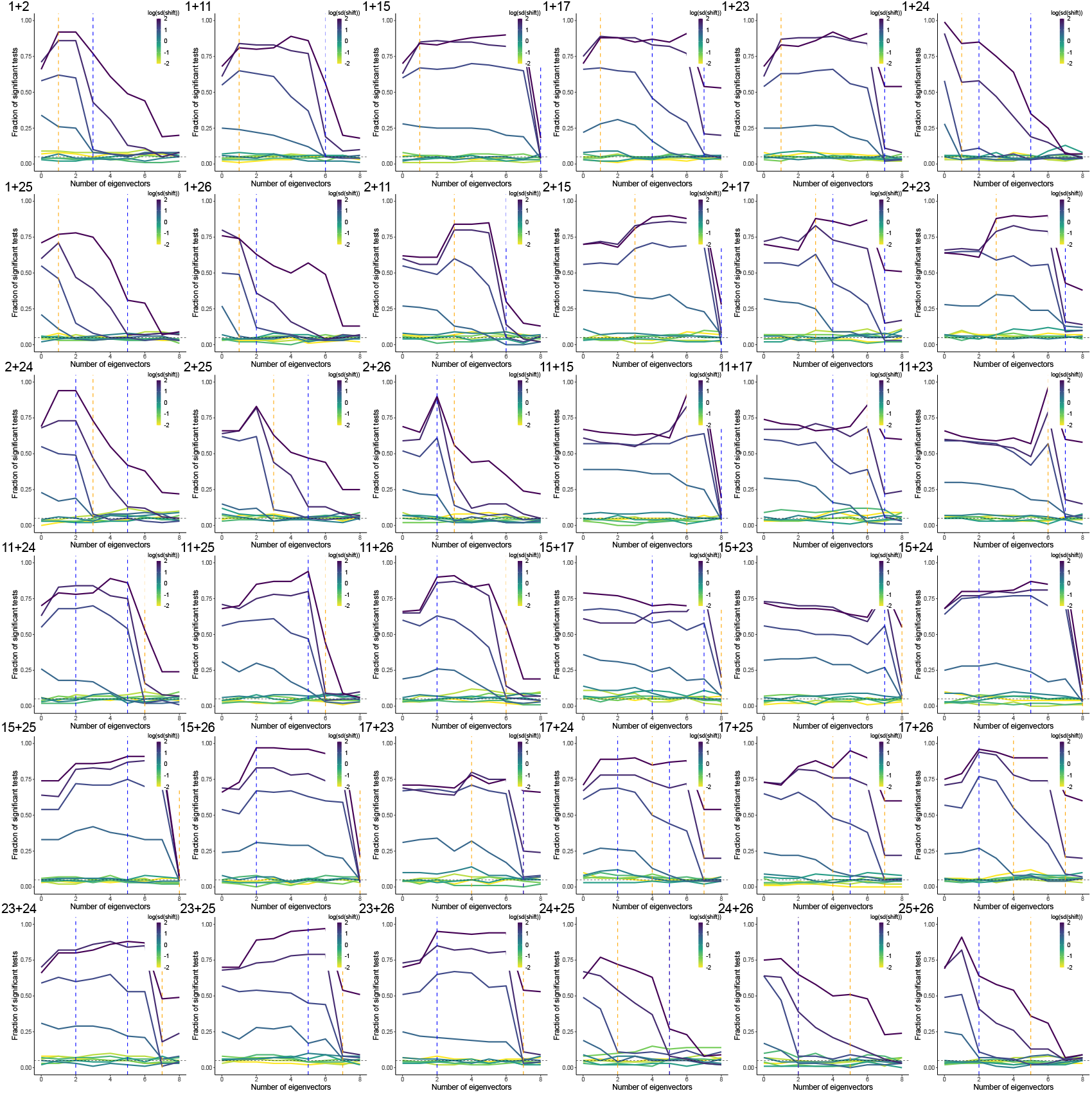
The performance of using eigenvectors in phylogenetic generalized least squares (PGLS) to remove phylogenetic confounding in a model with 16 tips related by a Yule tree with two non-Brownian shifts in both predictor and outcome variables. The shifts are simulated by adding a random normal variable with varying standard deviation to the species that are descendants of the branch. The label of each panel corresponds to the labels of the two branches where the shifts occur on the tree in Fig. S7. The horizontal axis shows the number of eigenvectors included as covariates, and the vertical axis shows the fraction of tests that would be significant at the 0.05 level. The vertical orange dashed lines show the dimensions to which the first branch concerned makes large contributions, and the vertical blue dashed lines show the dimensions to which the second branch concerned makes large contributions.

**Fig S12:**
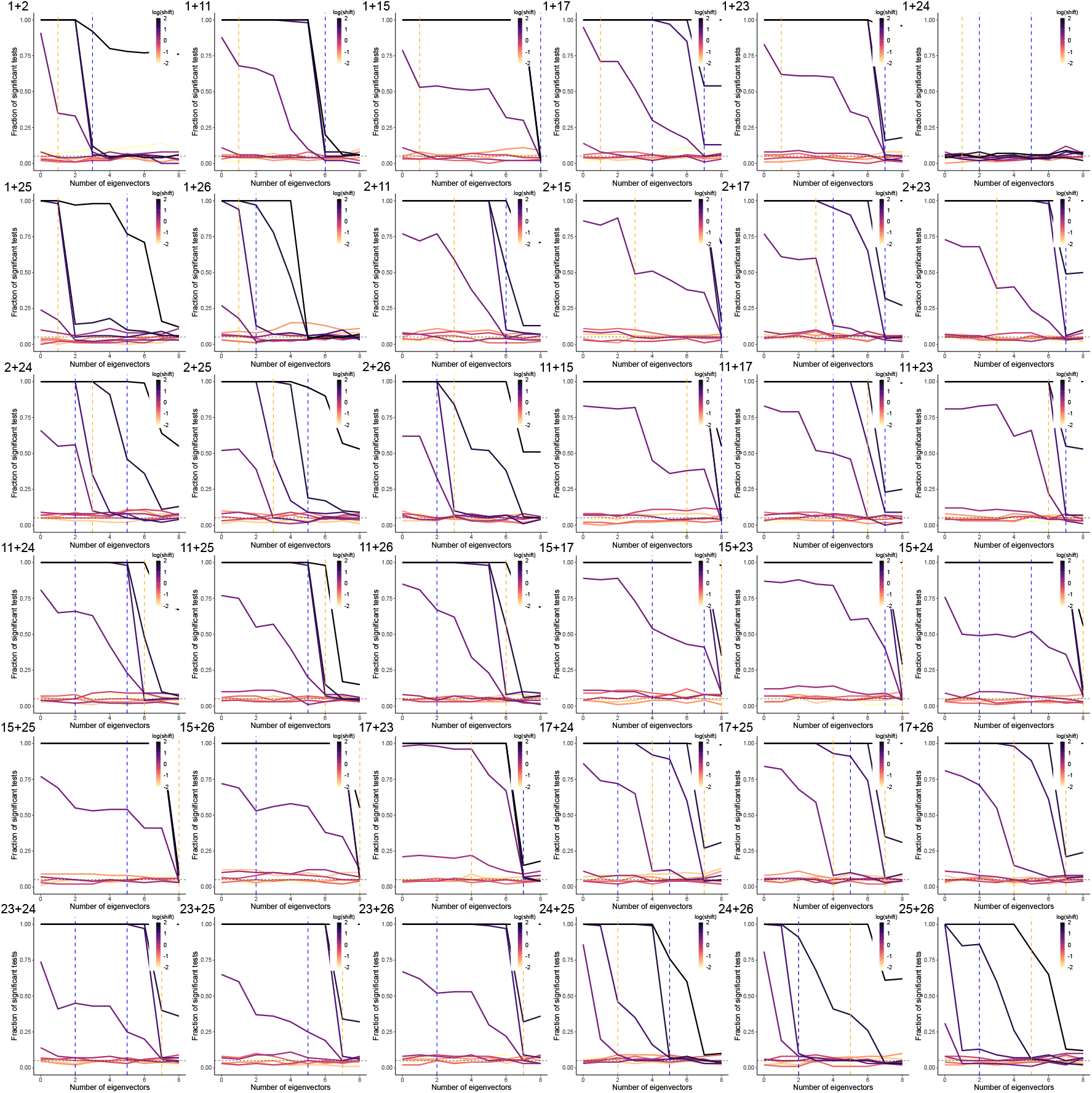
The performance of using eigenvectors in phylogenetic generalized least squares (PGLS) to remove phylogenetic confounding in a model with 16 tips related by a Yule tree with two non-Brownian shifts in both predictor and outcome variables. The shifts are simulated by adding a constant with varying size to the species that are descendants of the branch. The label of each panel corresponds to the labels of the two branches where the shifts occur on the tree in Fig. S7. The horizontal axis shows the number of eigenvectors included as covariates, and the vertical axis shows the fraction of tests that would be significant at the 0.05 level. The vertical orange dashed lines show the dimensions to which the first branch concerned makes large contributions, and the vertical blue dashed lines show the dimensions to which the second branch concerned makes large contributions.

**Fig S13:**
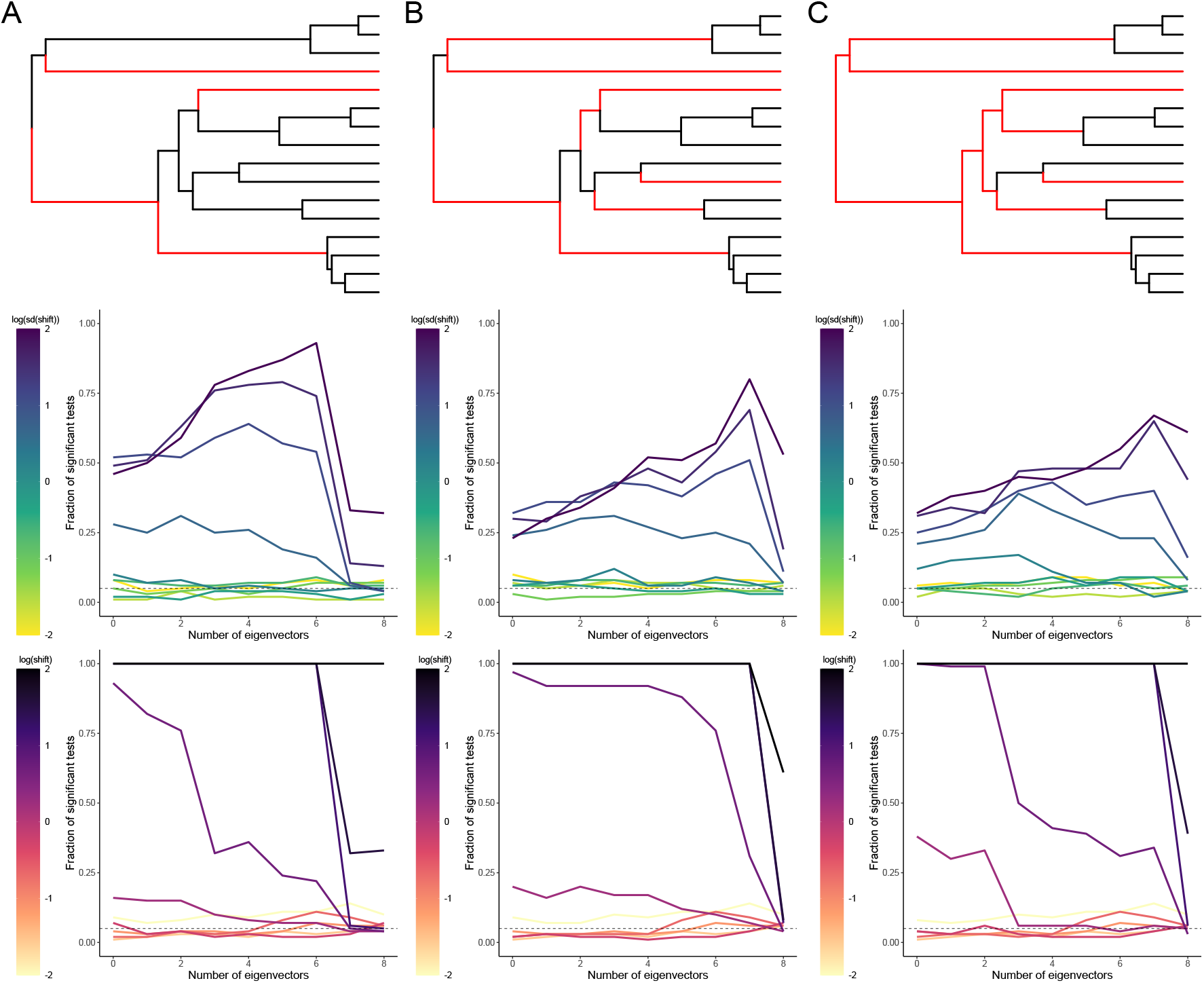
The performance of using eigenvectors in phylogenetic generalized least squares (PGLS) to remove phylogenetic confounding in a model with 16 tips related by a Yule tree with multiple non-Brownian shifts in both predictor and outcome variables. The top panels show the branches where the shifts occur in red. The middle panels show the performance of using eigenvectors in PGLS to remove phylogenetic confounding in the model, where the shifts are simulated by adding a normal random variable with varying standard deviation to the species that are descendants of the branch. The horizontal axis shows the number of eigenvectors included as covariates, and the vertical axis shows the fraction of tests that would be significant at the 0.05 level. The bottom panels show the performance of using eigenvectors in PGLS to remove phylogenetic confounding in the model, where the shifts are simulated by adding a constant of varying size to the species that are descendants of the branch. The horizontal axis shows the number of eigenvectors included as covariates, and the vertical axis shows the fraction of tests that would be significant at the 0.05 level. (A) There are 4 shifts on the tree. (B) There are 8 shifts on the tree. (C) There are 12 shifts on the tree.

**Fig S14:**
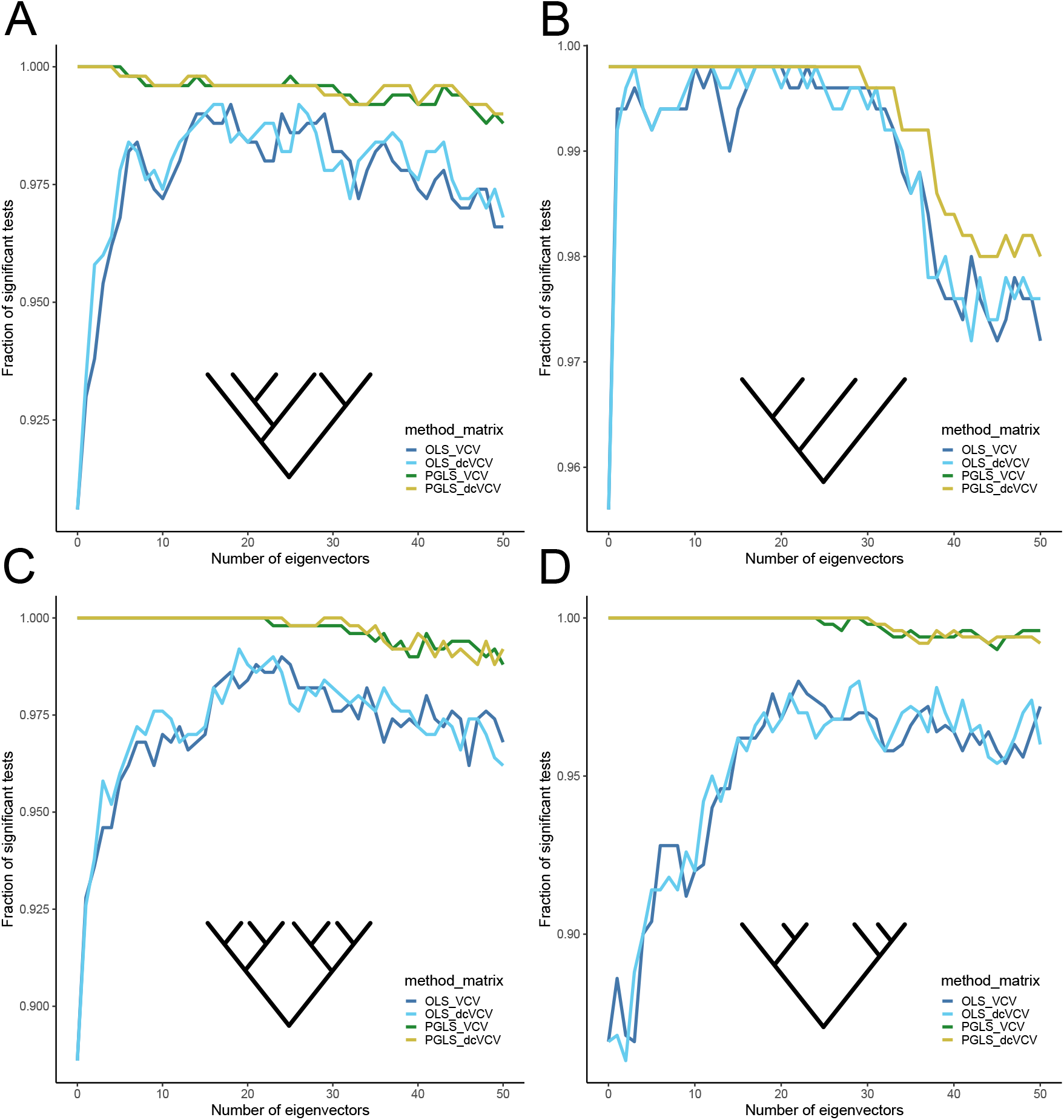
The performance of using eigenvectors of non-centered and double-centered phylogenetic variance - covariance matrices (*C* and 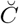) in ordinary least squares (OLS) and phylogenetic generalized least squares (PGLS) for different types of ultrametric trees when the true regression coefficient for *x, β*, is 0.5. (A) Performance of OLS and PGLS with eigenvectors of *C* and 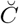 in models related by a Yule tree of 128 species. For simplicity, a Yule tree with only 6 tips is shown. The horizontal axis shows the number of eigenvectors included as covariates, and the vertical axis shows the fraction of tests that would be significant at the 0.05 level. (B) Performance of OLS and PGLS with eigenvectors of *C* and 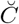 in models related by a caterpillar tree of 128 species. For simplicity, a caterpillar tree with only 4 tips is shown. The horizontal axis shows the number of eigenvectors included as covariates, and the vertical axis shows the fraction of tests that would be significant at the 0.05 level. The lines for PGLS_VCV and PGLS_dcVCV are overlapping. (C) Performance of OLS and PGLS with eigenvectors of *C* and 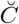 in models related by a fully balanced tree of 128 species. For simplicity, a fully balanced tree with only 8 tips is shown. The horizontal axis shows the number of eigenvectors included as covariates, and the vertical axis shows the fraction of tests that would be significant at the 0.05 level. (D) Performance of OLS and PGLS with eigenvectors of *C* and 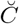 in models related by a coalescent tree of 128 species. For simplicity, a coalescent tree with only 6 tips is shown. The horizontal axis shows the number of eigenvectors included as covariates, and the vertical axis shows the fraction of tests that would be significant at the 0.05 level.

**Fig S15:**
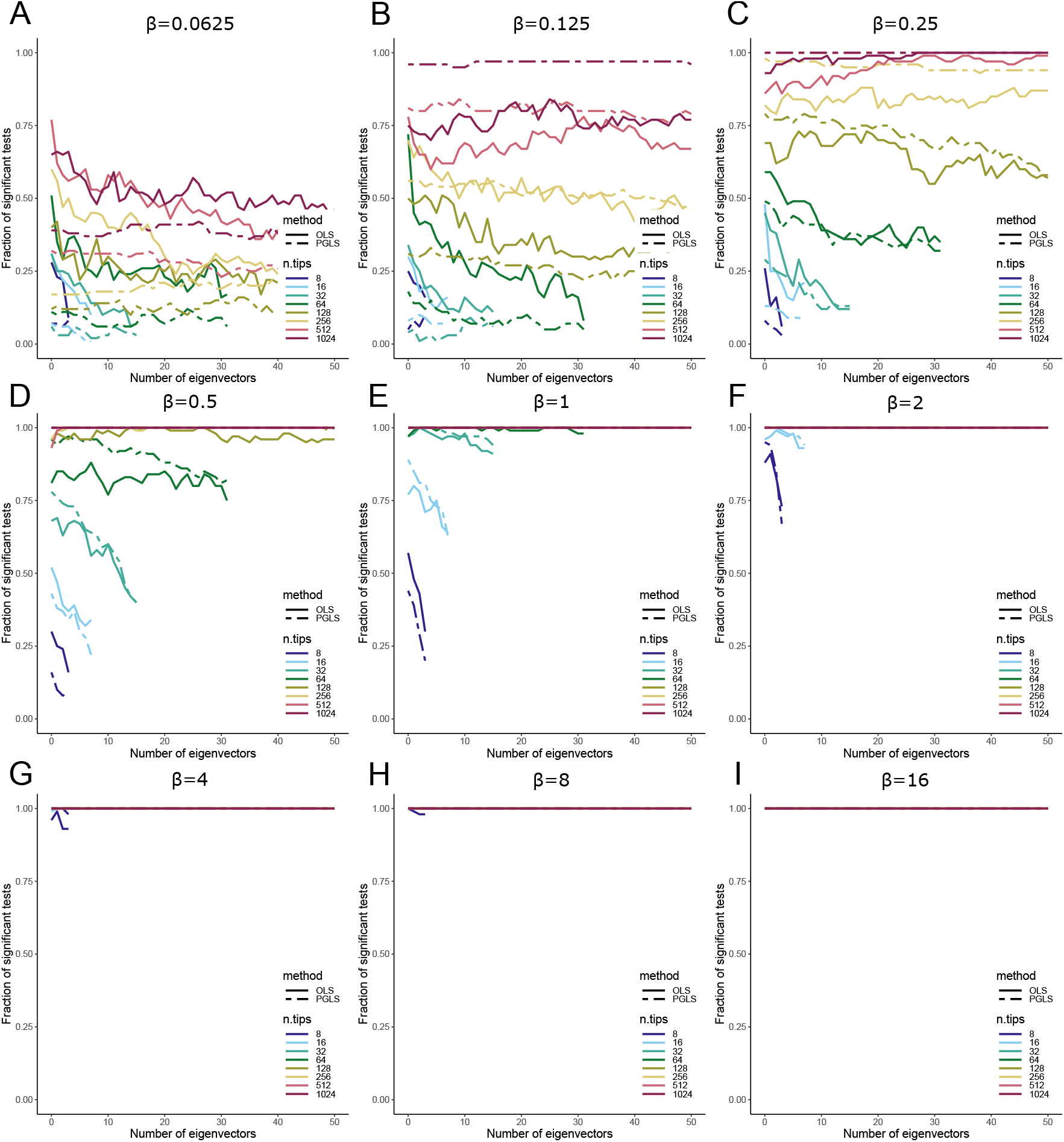
The recall after adding eigenvectors in ordinary least squares (OLS) and phylogenetic generalized least squares (PGLS) for Yule trees of varying numbers of species and different non-zero values of the true regression coefficient for *x, β*. The horizontal axes show the number of eigenvectors included as covariates, and the vertical axes show the fraction of tests that would be significant at the 0.05 level. (A) *β* = 0.0625. (B) *β* = 0.125. (C) *β* = 0.25. (D) *β* = 0.5. (E) *β* = 1. (F) *β* = 2. (G) *β* = 4. (H) *β* = 8. (I) *β* = 16.

**Fig S16:**
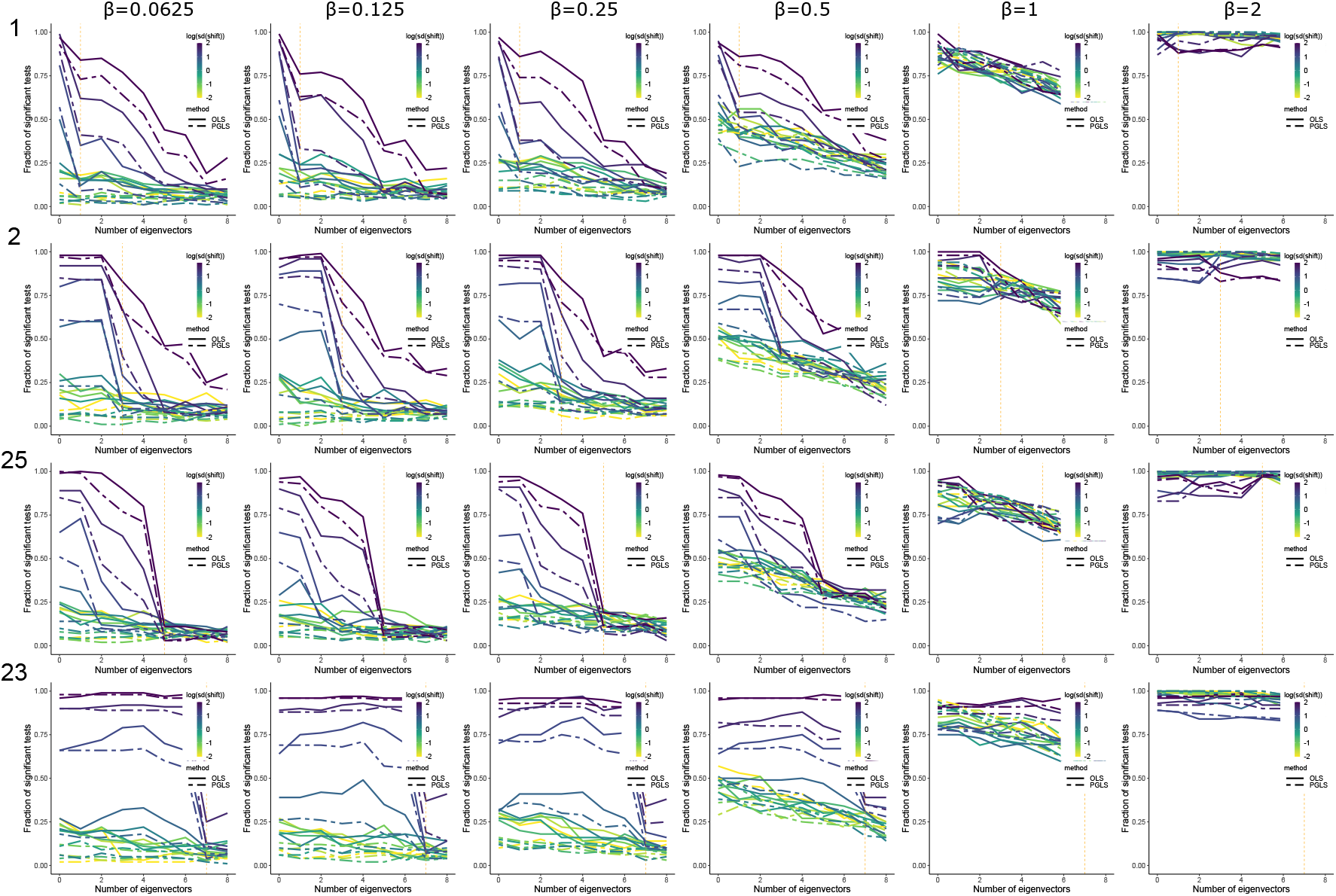
The recall after adding eigenvectors in ordinary least squares (OLS) and phylogenetic generalized least squares (PGLS) in a model with 16 tips related by a Yule tree with a non-Brownian shift in both predictor and outcome variables for different non-zero values of the true regression coefficient for *x, β*. The shift is simulated by adding a normal random variable with varying standard deviation to the species that are descendants of the branch. The horizontal axes show the number of eigenvectors included as covariates, and the vertical axes show the fraction of tests that would be significant at the 0.05 level. The numbers in front of each row indicate the labels of the branches where the shifts occur on the tree in Fig. S7, and each column corresponds to a different *β* value in 0.0625, 0.125, 0.25, 0.5, 1, and 2. The vertical orange dashed lines show the dimensions to which the branch concerned makes large contributions.

**Fig S17:**
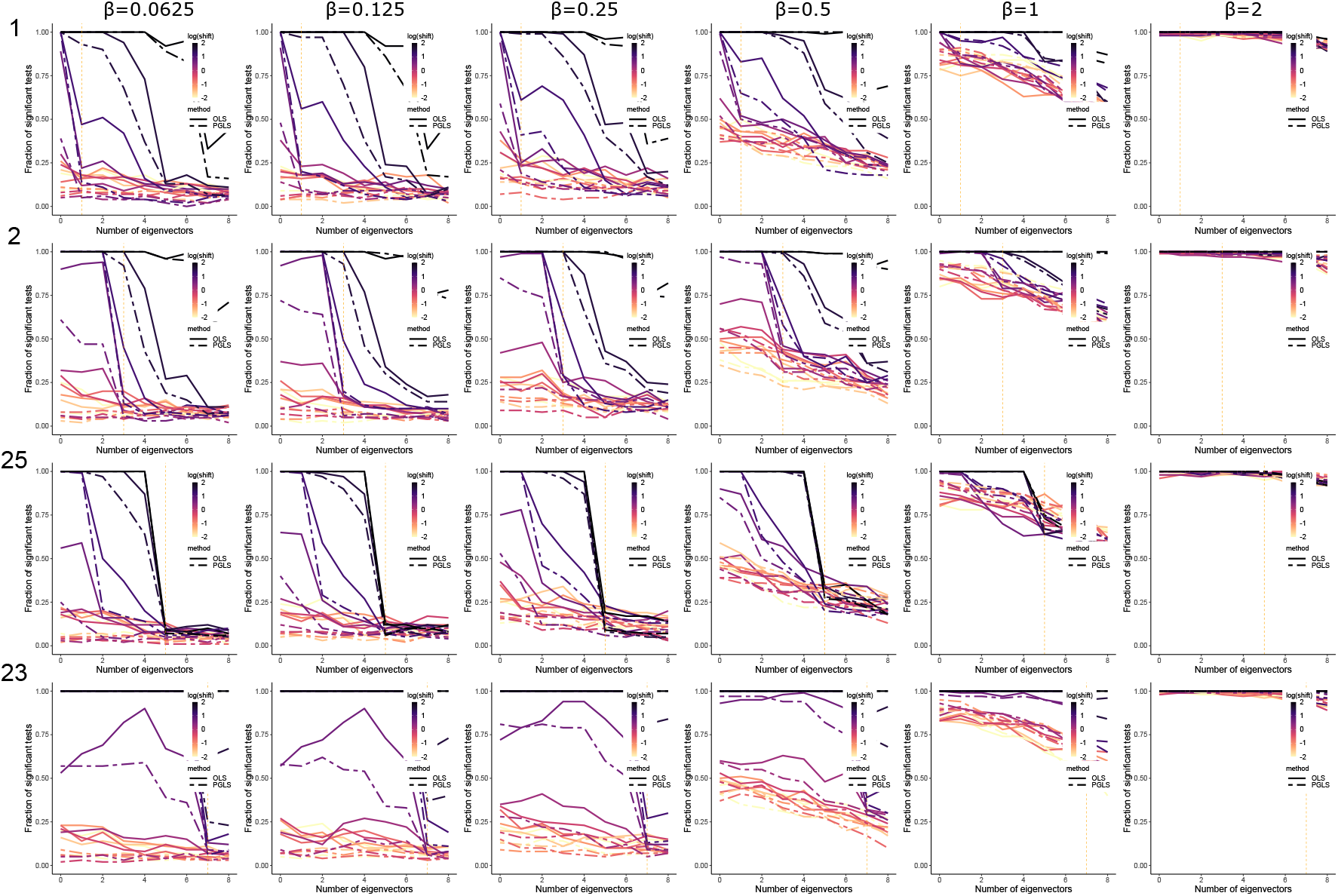
The recall after adding eigenvectors in ordinary least squares (OLS) and phylogenetic generalized least squares (PGLS) in a model with 16 tips related by a Yule tree with a non-Brownian shift in both predictor and outcome variables for different non-zero values of the true regression coefficient for *x, β*. The shift is simulated by adding a constant with varying size to the species that are descendants of the branch. The horizontal axes show the number of eigenvectors included as covariates, and the vertical axes show the fraction of tests that would be significant at the 0.05 level. The numbers in front of each row indicate the labels of the branches where the shifts occur on the tree in Fig. S7, and each column corresponds to a different *β* value in 0.0625, 0.125, 0.25, 0.5, 1, and 2. The vertical orange dashed lines show the dimensions to which the branch concerned makes large contributions.

